# Electrophysiological correlates of confidence differ across correct and erroneous perceptual decisions

**DOI:** 10.1101/2021.11.22.469610

**Authors:** Daniel Feuerriegel, Mackenzie Murphy, Alexandra Konski, Vinay Mepani, Jie Sun, Robert Hester, Stefan Bode

**Author notes:** **Corresponding Author:** Daniel Feuerriegel, Melbourne School of Psychological Sciences. Redmond Barry Building, Melbourne University, MN5N, Australia.

## Abstract

Every decision we make is accompanied by an estimate of the probability that our decision is accurate or appropriate. This probability estimate is termed our degree of decision confidence. Recent work has uncovered event-related potential (ERP) correlates of confidence both during decision formation and after a decision has been made. However, the interpretation of these findings is complicated by methodological issues related to ERP amplitude measurement that are prevalent across existing studies. To more accurately characterise the neural correlates of confidence, we presented participants with a difficult perceptual decision task that elicited a broad range of confidence ratings. We identified a frontal ERP component within an onset prior to the behavioural response, which exhibited more positive-going amplitudes in trials with higher confidence ratings. This frontal effect also biased measures of the centro-parietal positivity (CPP) component at parietal electrodes via volume conduction. Amplitudes of the error positivity (Pe) component that followed each decision were negatively associated with confidence for trials with decision errors, but not for trials with correct decisions, with Bayes factors providing moderate evidence for the null in the latter case. We provide evidence for both pre- and post-decisional neural correlates of decision confidence that are observed in trials with correct and erroneous decisions, respectively. Our findings suggest that certainty in having made a correct response is associated with frontal activity during decision formation, whereas certainty in having committed an error is instead associated with the post-decisional Pe component. These findings also highlight the possibility that some previously reported associations between decision confidence and CPP/Pe component amplitudes may have been a consequence of ERP amplitude measurement-related confounds. Re-analysis of existing datasets may be useful to test this hypothesis more directly.

**Highlights:**

– We mapped the event-related potential correlates of decision confidence
– A frontal component was associated with confidence during decision formation
– The error positivity component was associated with confidence in error trials
– The error positivity was not associated with confidence in correct response trials

## 1. Introduction

Every decision we make is accompanied by an estimate of the probability that our choice is accurate or appropriate for the task-at-hand. This probability estimate is known as our sense of decision confidence (Pouget et al., 2016). We can use our sense of confidence to decide whether to adjust our decision-making strategies in preparation for future events (van den Berg et al., 2016a; Desender et al., 2019a) and to rapidly correct decision errors when accuracy-related feedback is unavailable (Yeung & Summerfield, 2012). Computation of confidence is often conceptualised as a ‘second-order’ decision across a continuous dimension (e.g., ranging from ‘certainly wrong’ to ‘certainly correct’) that relates to a corresponding first-order decision (Yeung & Summerfield, 2012; Fleming & Daw, 2017). Within this framework, researchers have proposed two broad classes of theoretical models which delineate different sources of evidence that inform confidence judgments.

The first set of ‘decisional-locus’ models (as labelled in Yeung & Summerfield, 2012) assume that confidence judgments are based on information that directly relates to the first-order decision, such as the relative extent of evidence accumulated in favour of each choice alternative (Vickers, 1979; Vickers & Packer, 1982; Ratcliff & Starns, 2009; Kiani & Shadlen, 2009; Kiani et al., 2014). The other class of ‘postdecisional-locus’ models posit that confidence judgments are informed by processes which occur after the time of the initial decision, for example via post-decisional evidence accumulation (Rabbitt & Vyass, 1981; Pleskac & Busemeyer, 2010; Moran et al., 2015; van den Berg et al., 2016b; Desender et al., 2021a; Maniscalco et al., 2021) or motor action-related processes (e.g., Fleming & Daw, 2017; Turner et al., 2021). The main point of disagreement between these model classes relates to whether post-decisional processes are relevant to the computation of confidence (discussed in Moran et al., 2015; Fleming & Daw, 2017). For example, the decisional-locus model described in Vickers and Packer (1982) specifies no role of post-decisional evidence accumulation, whereas the model in Moran et al. (2015) specifies that post-decisional evidence accumulation is an important determinant of confidence judgments.

Because each account differs with respect to the timing of confidence-related computations relative to the first-order decision, electrophysiological measures with high temporal resolution, such as electroencephalography (EEG), have been used to identify neural correlates of decision confidence. This work has provided some support for both decisional and post-decisional locus models, however there are important methodological issues that limit the inferences we can draw from these studies.

### 1.1 Support for Decisional Locus Models

In line with predictions of decisional locus models, previous work has revealed that subjective and model-derived confidence ratings monotonically scale with the amplitude of the centro-parietal positivity (CPP) event-related potential (ERP) component (O’Connell et al., 2012) from around 300 ms after target stimulus onset (Squires et al., 1973; Gherman & Philiastides, 2015, 2018; Herding et al., 2019; Zarkewski et al., 2019; Rausch et al., 2020) or immediately preceding a keypress response used to report a decision (Philiastides et al., 2014). The CPP is thought to be analogous to the parietal P3 component in perceptual decision tasks (Twomey et al., 2015) and typically increases in amplitude to a fixed threshold around the time of a decision (O’Connell et al., 2012; Kelly & O’Connell, 2013; Twomey et al., 2015), closely resembling the accumulation-to-bound trajectories of decision variables in evidence accumulation models (Ratcliff, 1978; Ratcliff et al., 2016; Twomey et al., 2015; Kelly et al., 2021). These findings have been interpreted as reflecting higher levels of decision evidence accumulation in favour of the chosen option in trials with higher confidence ratings (e.g., Philiastides et al., 2014; Gherman & Philiastides, 2018). Consequently, this has been taken as support for the ‘balance of evidence hypothesis’ described in some decisional-locus models of confidence, which specifies that confidence indexes differences in the positions of racing accumulators in discrete choice tasks (Vickers, 1979; Vickers & Packer, 1982; Kiani & Shadlen, 2009). Here, we make the assumption that the CPP is time-locked to the response, in line with the original definition of this component (O’Connell et al., 2012). However, we note that in some studies the CPP and P3 are also considered to be stimulus-locked (e.g., Rausch et al., 2020).

The abovementioned studies have measured CPP/P3 amplitudes within a fixed time window relative to stimulus onset, except for Philiastides et al. (2014) which analysed model-derived (rather than self-reported) confidence ratings. Importantly, the CPP has been found to be tightly time-locked to the time of the keypress used to report a decision (e.g., O’Connell et al., 2012; van Vugt et al., 2019). Higher confidence ratings are typically given in trials with faster choice response times (RTs; e.g., Johnson, 1939; Vickers & Packer, 1982; Kiani et al., 2014), at least for a sizeable majority of individuals (for an analysis of 4,089 participants see Rahnev et al., 2020). In addition, participant-level RT distributions are strongly right-skewed, meaning that there is a larger amount of timing variability for relatively slower RTs. For example, there is typically a much larger range of RTs between the 70^th^ and 90^th^ percentiles of an RT distribution as compared to the range between the 10^th^ and 30^th^ percentiles. This means that, in many perceptual decision tasks, the CPP typically peaks within commonly-used stimulus-locked CPP/P3 amplitude measurement windows (e.g., 350-500ms in Rausch et al., 2020) in trials with faster RTs and higher confidence ratings (see Figure 1 of Kelly & O’Connell, 2013). In trials with slower RTs (and lower confidence ratings) the CPP is likely to peak later than these typical stimulus-locked measurement windows and will also show higher amounts of timing variability (i.e., temporal smearing, see Ouyang et al., 2015), producing apparently smaller stimulus-locked CPP amplitude measures in those trials. This, in turn, can artificially produce differences in stimulus-locked CPP amplitude measures across higher/lower confidence ratings in cases where there are no real differences during the pre-response time window in response-locked ERPs (e.g., O’Connell et al., 2012; Kelly & O’Connell, 2013). Consequently, based on our assumption that the CPP is a response-locked component, we believe it is important to measure CPP amplitudes using response-locked ERPs in addition to stimulus-locked measures, to ensure that effects on stimulus-locked ERPs are not simply by-products of RT differences across confidence rating conditions.

**Figure 1.**
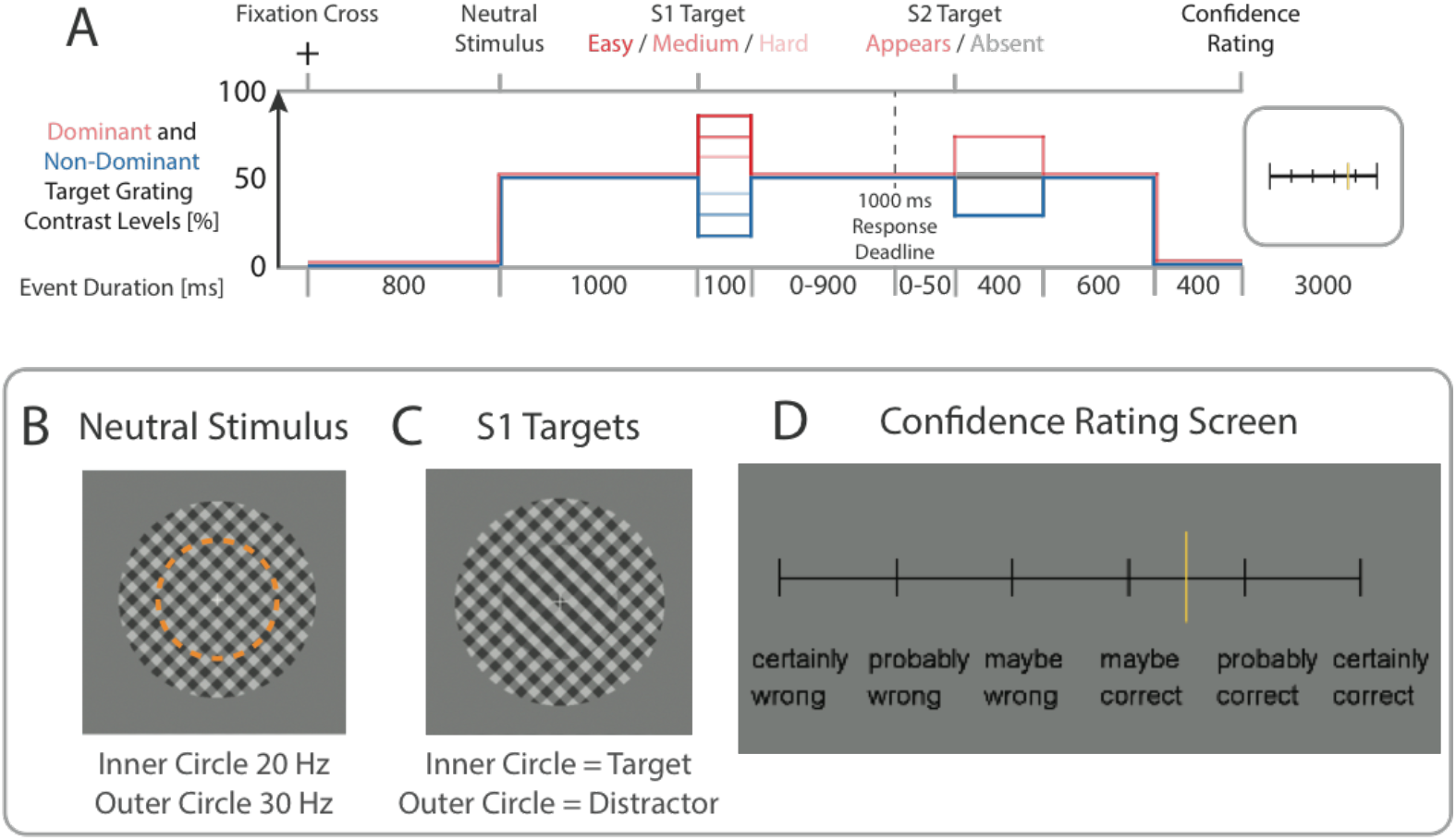
Trial structure and task. A) Each trial commenced with the presentation of a fixation cross. Following this, two circular apertures containing overlaid left- and right-tilted gratings were presented. Red and blue lines respectively depict the contrast levels of the dominant (i.e. higher contrast) and non-dominant (lower contrast) gratings within the target stimulus (inner circle) in each phase of the trial. Participants indicated which set of stripes in the S1 target stimulus was dominant (i.e. of higher contrast). In 50% of trials the S1 target was followed by a second S2 target that appeared within 50 ms of the response to the S1 target. Each S2 target contained a dominant grating at 75% contrast in the same direction as the preceding S1 target (S2 trials were not relevant to our research questions and were excluded from all confidence-related ERP analyses; see main text). Participants rated their decision confidence at the end of each trial. B) Example of the neutral stimulus. Gratings in the inner (target) and outer (distractor) circles contrast-reversed at 20 Hz and 30 Hz, respectively. The dashed orange line denotes the boundary between the inner (target) and outer (distractor) circles. Both the left- and right-tilted gratings within the distractor circle were kept constant at 50% contrast throughout the trial. C) Example of an S1 target stimulus with a dominant left-tilted grating. D) Confidence rating screen. Participants used their left and right index fingers to move the yellow cursor to their desired level of confidence.

The findings of a recent study (Kelly & O’Connell, 2013) also raise questions about whether these previously reported effects at centro-parietal electrodes actually reflect amplitude modulations of the CPP component. Kelly and O’Connell (2013) observed larger pre-response CPP amplitudes in conditions of higher stimulus discriminability (which often closely correlates with confidence). However, this effect was not observed after applying a current source density (CSD) transformation (Kayser & Tenke, 2006), which better isolates distinct cortical sources of EEG signals that are often conflated in analyses of standard ERPs. Kelly and O’Connell also identified a fronto-central component which exhibited more positive-going amplitudes in higher discriminability conditions, which appeared to bias CPP measurements at centro-parietal channels via volume conduction. CSD-transformations were not used in the primary data analyses of the other studies listed above, and so it remains to be verified if these findings reflect genuine modulations of the CPP, or contributions from other temporally-overlapping sources.

### 1.2 Support for Post-Decisional Locus Models

Studies supporting post-decisional locus models have described negative correlations between the amplitude of the error positivity (Pe) component (Falkenstein et al., 1991) and decision confidence ratings (Boldt & Yeung, 2015; Desender et al., 2019b). The Pe component occurs around 200-400 ms after the participant has formed a first-order decision and is measured at the same centro-parietal electrodes as the CPP component. A negative association between Pe amplitudes and confidence was first described by Boldt and Yeung (2015), who reported that Pe amplitudes were larger (i.e. more positive-going) when participants gave confidence ratings indicating that they had made an error, and also when they were less confident that they had made a correct decision. More specifically, they identified a monotonic relationship between confidence and Pe amplitude across the confidence spectrum ranging from ‘certainly wrong’ to ‘unsure’ to ‘certainly correct’. The Pe component was proposed to be a neural correlate of post-decisional evidence accumulation that is specifically framed in terms of detecting a response error, which in turn informs decision confidence judgements (Desender et al., 2021b; see also Murphy et al., 2015). Congruent with this notion, the Pe is also more positive-going when participants detect that they had committed a response error (e.g., Ridderinkhof et al., 2009; Steinhauser & Yeung, 2010; Wessel et al., 2011; Murphy et al., 2015).

Although Boldt and Yeung (2015) developed an innovative framework that attempted to unify confidence and error detection, there are also methodological issues that should be considered when interpreting their findings. Most importantly, for their key analyses they measured response-locked ERPs and used a pre-response baseline. Importantly, this pre-response baseline largely overlapped with a time window over which CPP/P3 amplitudes varied across confidence ratings, and the CPP and Pe were measured over similar sets of centro-parietal electrodes (their Figure 3B, see also Philiastides et al., 2014; Gherman & Philiastides, 2015, 2018). In such cases, any systematic differences across conditions during the pre-response baseline period will lead to spurious differences in post-response ERPs (Luck, 2014). In the case of Boldt and Yeung (2015), their baseline subtraction procedure would have artificially inflated Pe amplitudes in trials with lower pre-response CPP amplitudes, such as trials with lower confidence ratings or response errors (e.g., von Lautz et al., 2019). This issue also applies to a subsequent study that replicated this effect (Desender et al., 2019b). Therefore, it remains to be verified whether Pe amplitudes do monotonically scale with decision confidence ratings in perceptual decision tasks (e.g., as claimed by Desender et al., 2021b).

### 1.3 The Present Study

To more accurately characterise associations between decision confidence and CPP and Pe component amplitudes, we presented a difficult perceptual discrimination task and required participants to give confidence ratings in each trial. To better understand the sources of pre-decisional ERP correlates of confidence, we assessed the effects of applying CSD transforms to our data in line with Kelly and O’Connell (2013). We expected to find response-locked CPP amplitudes to be positively correlated with confidence in trials with correct responses (as reported in Philiastides et al., 2014; Rausch et al., 2020), however we were agnostic about whether this effect would remain once a CSD transform had been applied. We also investigated the effects of using target stimulus- and response-locked baselines on associations between confidence and Pe amplitudes. We predicted that the associations between decision confidence and Pe amplitudes reported in Boldt and Yeung (2015) would not be replicated when using a pre-stimulus ERP baseline, but would be artificially produced when using a pre-response baseline.

## 2. Method

### 2.1. Participants

Thirty-five people (20 female, 15 male, aged 18-36 years, M = 24.7, SD = 4.8) participated in this experiment. Participants were right-handed, fluent in English and had normal or corrected-to-normal vision. Four participants were excluded due to near-chance task performance (i.e., accuracy below 55% for any of the three stimulus discriminability conditions). One additional participant was excluded due to excessively noisy EEG data. One participant was excluded because they were unable to complete the task, leaving 29 participants for both behavioural and EEG data analyses (17 female, 12 male, aged 18-36 years, M = 25.0, SD = 4.9). This study was approved by the Human Ethics Committee of the Melbourne School of Psychological Sciences (ID 1750871).

### 2.2 Stimuli

Stimuli were presented using a gamma-corrected 24” Benq RL2455HM LCD monitor with a refresh rate of 60 Hz. Stimuli were presented using functions from MATLAB (Mathworks) and PsychToolbox (Brainard, 1997; Kleiner et al., 2007). Code used for stimulus presentation will be available at https://osf.io/gazx2/ at the time of publication.

The critical stimuli consisted of two overlaid diagonal gratings within a circular aperture, presented against a grey background (similar to Steinemann et al., 2018; Feuerriegel et al., 2021a). The two gratings were oriented 45° to the left and right of vertical, respectively. The circular aperture was divided into two concentric circles: an inner circle (target stimulus) and an outer circle (distractor) with radii of 2.86° and 5.31° of visual angle (Figure 1B). The inner circle contrast-reversed at a rate of 20 Hz; the outer circle contrast-reversed at a rate of 30 Hz (which allowed for frequency tagging of target and distractor stimuli in analyses of steady state visual evoked potentials, however these signals were not relevant to the research question of this paper).

### 2.3 Procedure

Participants sat 80 cm from the monitor in a darkened room and were asked to fixate on a central cross throughout all trials. The trial structure is depicted in Figure 1A. In each trial, a white fixation cross appeared for 800 ms. Following this, both left- and right-tilted gratings within each circle increased from 0% to 50% contrast. Both gratings remained at 50% contrast for a further 1,000 ms, during which the contrast levels of both gratings were identical (i.e. the stimulus was “neutral”, see Figure 1B).

Immediately after this neutral stimulus period, one of the gratings within the inner circle increased in contrast and the other decreased in contrast (see Figure 1C; labelled the “S1” target). This contrast difference persisted for 100 ms, after which the neutral stimulus was presented again. Participants indicated which grating within the inner circle (i.e. left-tilted or right-tilted) was dominant (i.e. of higher contrast) by pressing keys on a TESORO Tizona Numpad (1,000 Hz polling rate) using their left and right index fingers. Participants were required to respond within 1,000 ms of S1 target onset. If responses were made prior to the S1 target onset, or after the 1,000 ms S1 response deadline, then “Too Early” or “Too Slow” feedback appeared, respectively. Feedback signalling the accuracy of the decision was not provided.

Relative contrast levels of the dominant and non-dominant gratings varied throughout the experiment according to an accelerated stochastic approximation (ASA) staircase procedure (Kesten, 1958; initial step size = 0.25, minimum contrast level = 0.51, maximum contrast level = 0.95). Rather than using an up-down staircase procedure that only converges on a small number of accuracy ratings, this ASA procedure was used because it quickly converges to pre-specified accuracy targets (Lu & Dosher, 2013). We used three different staircases (interleaved across trials) that were designed to converge to accuracy levels of 60%, 75% and 90%. The staircase procedure was employed continuously throughout the experiment to account for any improvements in task performance that can occur across the first few blocks of an experiment, and to provide a wide range of stimulus contrast values (and a wide range of confidence ratings). In each trial, staircase 1, 2 or 3 was pseudorandomly selected to determine the contrast level of the target stimulus. Equal numbers of each staircase condition were presented within each block, and across the experiment. Both the left- and right-tilted gratings within the outer distractor circle were kept constant at 50% contrast throughout each trial. The purpose of presenting the distractors was to increase the difficulty of the task by encircling the target with a dynamically contrast-reversing neutral stimulus.

Following the response to the S1 target, the neutral stimulus was presented for a further 1,000 ms in 50% of trials. In the other 50% of trials a second S2 target was presented at the time of the next inner circle (target) contrast reversal after the response (i.e. within 50 ms following the response), whereby the dominant S1 grating in the inner circle was again presented at 75% contrast, and the non-dominant S1 grating at 25% contrast. This second target was presented for 400 ms, after which a neutral stimulus was presented for 600 ms. Note that the dominant grating orientation was always consistent across the first and second targets, meaning that the second target was informative as to the correct response in that trial. Participants were instructed not to respond to the second target but were advised that the information conveyed by this stimulus would be useful in forming their confidence judgements in the trials in which it appeared. These S2 targets were originally included to investigate the neural correlates of decision updating that occurs when additional information is provided after making a perceptual decision (similar to Fleming et al., 2018), and the respective analyses will be part of a separate publication. Importantly, none of the analyses testing for associations between ERPs and confidence presented here included the trials in which the S2 target appeared. Numbers of trials with and without the second target were balanced within each staircase condition. In all trials the grating stimuli were then replaced with a blank screen with a fixation cross for 400 ms.

Participants then rated their confidence in their decision on a continuous scale (ranging from -100 to 100) with equal intervals between the labels ‘Certainly Wrong’, ‘Probably Wrong’, ‘Maybe Wrong’, ‘Maybe Correct’, ‘Probably Correct’ and ‘Certainly Correct’ (Figure 1D). Please note that these labels were indicators to guide selection of a continuously-valued confidence response, but not discrete rating choice options. The zero value was the midpoint of the scale, indicating maximal uncertainty as to whether a correct response or an error had occurred. To provide confidence ratings, participants held down the left and right response keys to move a vertical bar to the left or right on the scale. To discourage premature preparation of motor responses associated with specific confidence ratings, the vertical bar was initially placed in a random location on the scale in each trial. The confidence rating scale was presented for 3,000 ms. The location of the bar at the end of this period constituted the confidence rating for that trial.

To encourage participants to perform the task with maximum accuracy and make confidence judgements that reflected their true degree of belief in the correctness of their choices, we implemented a points system based on both task performance and the correspondence between participants’ confidence ratings and their objective accuracy within each trial (as done by Fleming et al., 2018). Participants were awarded 10 points if they made a correct decision in each trial. No points were lost if the decision was incorrect. Trials with more than one response, responses prior to S1 target onset, or no response within the 1,000 ms deadline, resulted in a loss of 50 points.

Participants could gain or lose up to an additional 50 points by making an accurate confidence rating regarding their response to the S1 target. As the confidence responses were graded, the most extreme confidence ratings were associated with the highest number of points wagered. An accurate confidence rating resulted in a gain between 1-50 points; a confidence rating in the incorrect direction resulted in a loss of between 1-50 points. For example, if a participant made a correct response and moved their rating bar halfway toward ‘certainly correct’ from the midpoint, they would win 25 points. In order to encourage optimal performance, participants were told they could earn between 20-25 AUD based on how many points they accumulated, with 1 AUD awarded for every 5,000 points obtained (maximum possible score = 28,800). However, all participants were actually reimbursed 25 AUD at the end of the experiment.

The experiment consisted of eight blocks, each containing 60 trials (total number of trials = 480). This included 240 trials where the S2 target appeared, and 240 trials in which it did not appear. Participants could take self-paced breaks between each block (minimum break length = 30 seconds). Prior to the experiment participants completed a brief practice block of 20 trials. During this practice phase, participants received feedback at the end of each trial as to whether their response was correct or an error. Participants were allowed to repeat the practice block until both they and the experimenter were confident that they understood the task.

### 2.4. Analyses of Accuracy, Response Times and Confidence Ratings

Code used for all behavioural and EEG data analyses will be available at https://osf.io/gazx2/ at the time of publication. Trials with responses slower than the response deadline or earlier than S1 target stimulus onset were removed from the dataset. Only trials with correct or erroneous responses and response times (RTs) of >100 ms were included for analyses of RTs. For analyses of accuracy and RTs we included trials in which the S2 target appeared because this target was presented after the time of the response to the S1 target and so could not influence these measures. For analyses of confidence ratings, we excluded trials where the S2 target appeared. We modelled proportions of correct responses using generalised linear mixed effects logistic regressions (binomial family) as implemented in the R package lme4 (Bates et al., 2015). We modelled RTs using generalised linear mixed effects regressions (Gamma family, identity link function) as recommended by Lo and Andrews (2015). We modelled confidence ratings using linear mixed effects models (Gaussian family) as done by Fleming et al. (2018).

To test for effects of each factor of interest on measures of accuracy, RTs and confidence ratings, we compared models with and without that fixed effect of interest using likelihood ratio tests. For each comparison, both models included identical random effects structures, including random intercepts by participant. The fixed effect of interest in all analyses was target discriminability (i.e. contrast level). The fixed effect of correct/error trial outcome was included in all models for RT analyses. Random slopes were also included for effects of target discriminability (Accuracy and RT analyses) and trial outcome (RT analyses) as these models converged successfully. Models of confidence ratings were fit to correct and error trials separately (as done by Fleming et al., 2018). The structure of each model and the coefficients of each fitted model are detailed in the Supplementary Material.

### 2.5. EEG Data Acquisition and Processing

We recorded EEG at a sampling rate of 512 Hz from 64 active electrodes using a Biosemi Active Two system (Biosemi). Recordings were grounded using common mode sense and driven right leg electrodes (http://www.biosemi.com/faq/cms&drl.htm). We added six additional channels: two electrodes placed 1 cm from the outer canthi of each eye, and electrodes placed above and below the center of each eye.

We processed EEG data using EEGLab v13.4.4b (Delorme & Makeig, 2004). All data processing and analysis code and data will be available at https://osf.io/gazx2/ at the time of publication. First, we identified excessively noisy channels by visual inspection (median number of bad channels = 2, range 0-8) and excluded these from average reference calculations and Independent Components Analysis (ICA). Sections with large artefacts were also manually identified and removed. We re-referenced the data to the average of all channels, low-pass filtered the data at 40 Hz (EEGLab Basic Finite Impulse Response Filter New, default settings), and removed one extra channel (AFz) to correct for the rank deficiency caused by the average reference. We processed a copy of this dataset in the same way and additionally applied a 0.1 Hz high-pass filter (EEGLab Basic FIR Filter New, default settings) to improve stationarity for the ICA. ICA was performed on the high-pass filtered dataset (RunICA extended algorithm, Jung et al., 2000). We then copied the independent component information to the non high-pass filtered dataset (e.g., as done by Feuerriegel et al., 2018). Independent components generated by blinks and saccades were identified and removed according to guidelines in Chaumon et al. (2015). After ICA we interpolated any excessively noisy channels and AFz using the cleaned data (spherical spline interpolation). EEG data were then high-pass filtered at 0.1 Hz (EEGLab Basic Finite Impulse Response Filter New, default settings).

The resulting data were segmented from -3,200 ms to 4,000 ms relative to S1 target onset, and were baseline-corrected using the -200 to 0 ms pre-target interval (note that, for some analyses described below, ERPs were baseline-corrected relative to a pre-response baseline at a later step). These long epochs were derived to also allow for analyses of SSVEPs and time-frequency data, which are not relevant to the research questions here. Epochs containing amplitudes exceeding ±200 µV at any scalp channels between -500 ms and 2,500 ms from S1 target onset were rejected (mean trials retained = 405 out of 480, range 289-456). This long time window was used to ensure that the same epochs were included for analyses of both stimulus- and response-locked ERPs. Numbers of retained epochs by condition are displayed in Supplementary Tables S1, S2). From the resulting epoched data, we then derived stimulus-locked epochs using the interval from -500 ms to 1,000 ms relative to the S1 target onset. We also derived response-locked epochs using the interval from -1,500 ms to -1,500 ms relative to the time of the next inner circle contrast reversal after the response to the S1 target. Because the gratings in the inner circle contrast-reversed at a rate of 20 Hz (i.e., every 50 ms), the time point for deriving response-locked epochs always occurred within a very short latency (0-50 ms) following the keypress response. This epoching method was used to align the timing of target stimulus contrast-reversals across conditions, so that there would be no systematic discrepancies in the timing of visual evoked responses associated with these reversals. This likely resulted in a small amount of temporal smearing of response-locked ERP components, the extent of which is smaller than the width of the time windows used to measure the ERP components of interest. We also note that, although the inner circle is termed the target stimulus, the inner circle actually contained a neutral stimulus (i.e., left- and right-tilted gratings at equal contrast) at the time of the keypress response in each trial. Please also note that the derived epochs extended beyond the time windows used for ERP analyses to also allow analyses of steady state visual evoked potentials (SSVEPs) and time-frequency power measures. However, such measures were not directly relevant to the research questions of this paper and are not reported here.

### 2.6. ERP Component Amplitude Analyses

We measured mean ERP amplitudes of the pre-response CPP between -130 to -70 ms relative to the response at parietal electrodes Pz, P1, P2, CPz, and POz (same time window as Steinemann et al., 2018; Feuerriegel et al., 2021a). For these analyses we used a pre-stimulus baseline (i.e., the -200 to 0 ms pre-target interval). To link our results to previous work using stimulus-locked CPP measures (e.g., Gherman & Philiastides, 2018; Rausch et al., 2020) we also measured the CPP as the mean amplitude between 350-500 ms from S1 target onset (as done by Rausch et al., 2020). For the reasons described in section 1.1 we do not focus on the results of these analyses in our paper. However, we acknowledge that these results may be interesting to those who assume that the CPP is best understood as a component that is time-locked to the stimulus rather than the response.

For analyses of the Pe component we first derived single-trial ERPs using the pre-stimulus baseline described above. To more directly compare our results with those of Boldt and Yeung (2015), we additionally ran the same set of Pe mean amplitude analyses using a -100 to 0 ms pre-response baseline. We measured Pe amplitudes as the mean amplitude between 200-350 ms relative to the response, at the same set of parietal electrodes as for the CPP (same time window as Nieuwenhuis et al 2001; Di Gregorio et al., 2018, and similar to the 250-350 ms window in Boldt & Yeung, 2015).

For analyses of the CPP component we compared correct and erroneous responses using paired-samples frequentist and Bayesian t-tests as implemented in JASP v0.9.1 (JASP Core Team; Cauchy prior distribution, width 0.707, default settings). We additionally fitted linear regression models using MATLAB to predict mean amplitudes based on confidence ratings. This was done separately for analyses of trials with correct responses and trials with errors. The resulting Beta coefficients (slopes) were tested at the group-level using one-sample frequentist and Bayesian t-tests (as done by Feuerriegel et al., 2021b). For analyses of the CPP the correct/error comparison included both trials whereby the S2 target did and did not appear, as the stimulation conditions were not systematically different until the time of the response. For analyses including confidence ratings, only trials whereby the S2 target did not appear were included. As described above, this is because the appearance of this informative, easily-discriminable S2 target systematically biased confidence ratings toward the extremes of the rating scale.

Analyses of the Pe component used data from trials in which the S2 target did not appear. Paired-samples t-tests were conducted to compare Pe amplitudes across trials with correct and erroneous responses. Within-subject regressions and group-level t-tests were performed using the predictor of confidence as described above.

We also performed complimentary, post hoc regression analyses using restricted ranges of confidence ratings, including the range from “unsure” (0) to “certainly correct” (100; indexing participants’ certainty that they had made a correct response) and, in separate analyses the range from “certainly wrong” (−100) to “unsure” (indexing participants’ certainty that they had made an error). For these analyses, we included both trials with correct responses and errors. We only included trials in which the S2 target did not appear. The results for each ERP component are included in the Supplementary Material.

### 2.7. Current Source Density Transformation

Based on observed positive associations between CPP amplitudes and confidence, we repeated the CPP analyses using CSD-transformed data estimated using the CSD Toolbox (Kayser & Tenke, 2006; m-constant = 4, λ = 0.00001). For analyses of CSD-transformed data we selected slightly different sets of electrodes to better isolate localised effects that become apparent when using this data transformation. We measured the CPP at CPz and Pz, consistent with electrodes used in previous work (e.g., O’Connell et al., 2012; Kelly & O’Connell, 2013; Steinemann et al., 2018; Feuerriegel et al., 2021a). We selected these electrodes because the CPP component shows a very focal distribution over Pz and CPz in CSD-transformed data, with amplitudes that can be markedly diminished at neighbouring channels (e.g., Kelly & O’Connell, 2013; Murphy et al., 2015; Feuerriegel et al., 2021). Based on correlations between stimulus discriminability (which correlates with confidence) and fronto-central electrode amplitudes reported by Kelly and O’Connell (2013), we additionally measured amplitudes of a frontal component at channel FCz during the time window used to measure the CPP. Note that we did not analyse Pe amplitudes using CSD-transformed data because our comparison to the findings of Boldt and Yeung (2015) relies on non-transformed data only. We did not have any *a priori* hypotheses about the influence of CSD transformations during this time window.

## 3. Results

### 3.1. Task Performance Results

The interleaved staircase procedure had the intended effects on measures of accuracy, RT and decision confidence (plotted in Figure 2). Group-averaged mean contrast levels of the dominant S1 gratings were 71%, 80% and 91% for staircases 1 (low discriminability), 2 (medium discriminability) and 3 (high discriminability). Group mean contrast levels by staircase across the course of the experiment are displayed in Supplementary Figure S1. Responses prior to S1 target onset occurred in only 1% of trials within each staircase on average, and responses slower than the deadline occurred in 7%, 3% and 2% of trials for staircases 1, 2, and 3. Participants accrued on average 17,865 points out of a total of 28,800 points (SD = 3,744, range = 7,865-23,086) by the end of the experiment.

**Figure 2.**
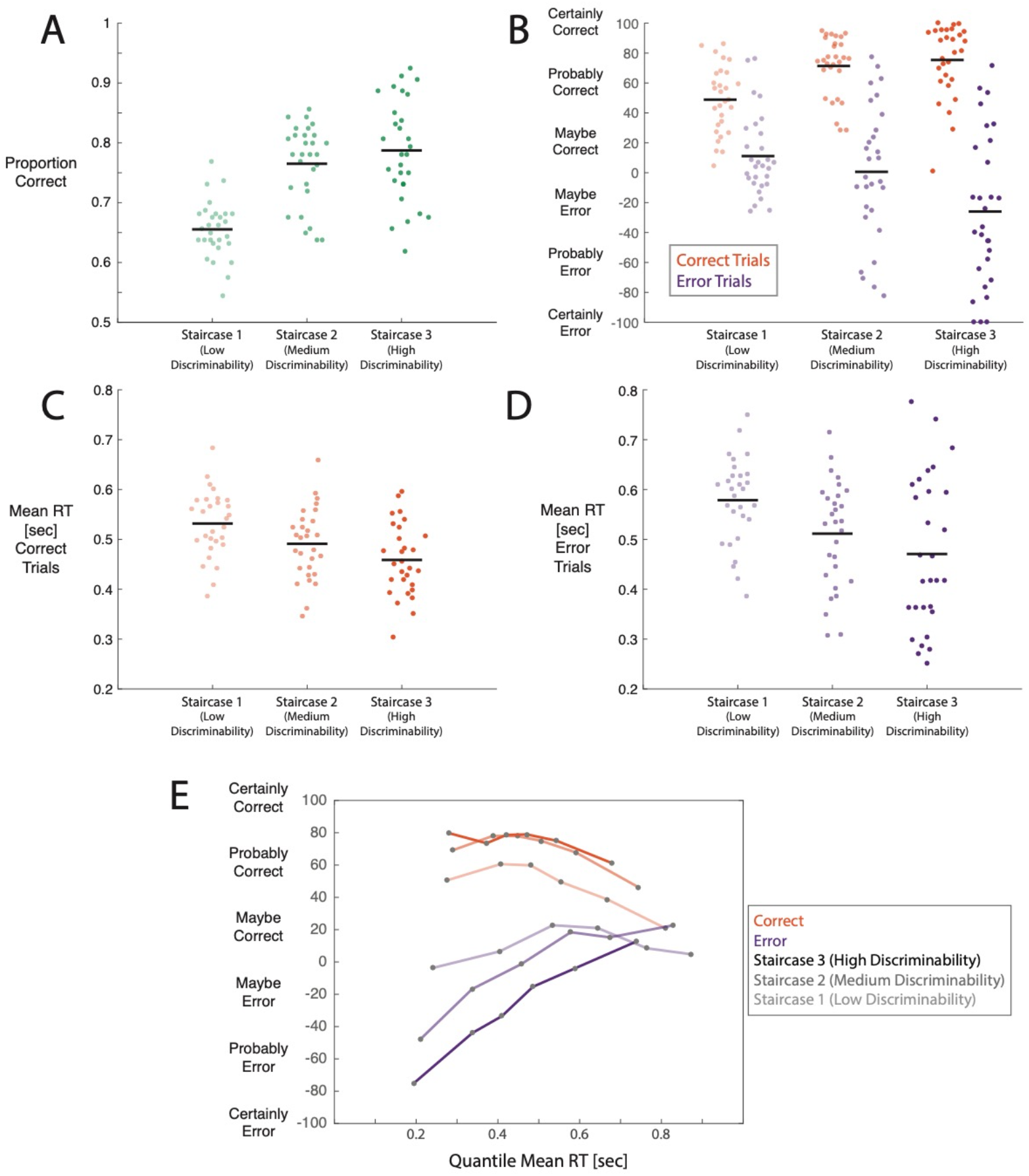
Task performance plotted by staircase. A) Proportions of correct responses (i.e., accuracy scores). B) Mean confidence ratings for trials with correct and erroneous responses, for trials whereby the S2 target did not appear. C) Mean RTs for trials with correct responses. D) Mean RTs for trials with erroneous responses. For A-D, black lines denote group mean values and dots represent individual participant values. E) Mean confidence ratings by RT quantile for correct and error responses in each staircase condition where the S2 target did not appear. Boundaries between quantiles were set at the 10, 30, 50, 70 and 90^th^ percentiles of the RT distributions.

Increasing target discriminability led to higher accuracy (likelihood ratio test χ^2^ (1) = 16.31, p < .001), faster RTs (χ^2^ (1) = 26.79, p < .001) and higher levels of confidence (χ^2^ (1) = 303.47, p < .001) in trials with correct responses (Figure 2A-D). For trials with errors, higher stimulus discriminability was instead associated with lower confidence ratings (Figure 2B, χ^2^ (1) = 44.68, p < .001, as also reported in Sanders et al., 2016; Desender et al., 2019a; Turner et al., 2021; but see Kiani et al., 2014; Rausch et al., 2018, 2020 for opposite patterns of effects). RTs were slower in trials with errors compared to correct responses (RT model fixed effect t = -3.07, p = .002). Tables of model coefficients are included in the Supplementary Material.

To display qualitative relationships between confidence, response speed and target discriminability, Figure 2E shows mean confidence ratings by RT quantile and staircase condition, for both correct and error responses. For trials with correct responses, confidence was on average lower in trials with slower RTs in all staircase conditions. In trials with decision errors, confidence increased toward the scale midpoint (i.e., unsure whether correct or an error) in trials with slower RTs. RT histograms for different confidence rating bands are displayed in Supplementary Figure S2.

Here we note that, although the patterns of mean confidence judgments in our group-level results are consistent with those observed in previous studies, there was substantial inter-individual variation in the distributions of confidence ratings in correct and error trials. For example, some participants used the entire range of the confidence scale, whereas others concentrated their responses within a narrower band (for distributions of confidence ratings by participant see Supplementary Figure S3). Accordingly, to better display these results, we have plotted ERPs corresponding to trials with lower and higher confidence ratings (determined using median splits within correct/error conditions) rather than by confidence rating band. Results of the regression analyses predicting ERP component mean amplitudes using confidence ratings are plotted in Supplementary Figure S4. Importantly, the results of the regression analyses do not differ from the patterns of effects indicated by the ERP plots.

### 3.2. CPP Amplitudes

CPP amplitudes (that were not CSD transformed, and measured using pre-stimulus baseline-corrected ERPs) were positively associated with confidence ratings for both correct responses and errors. Response-locked ERP waveforms are plotted in Figure 3. CPP amplitudes were larger (i.e., more positive-going) when preceding correct responses compared to errors (t(28) = 7.72, p < .001, BF_10_ > 650,000, Figure 3A). CPP amplitudes were positively associated with decision confidence ratings both in trials with correct responses (t(28) = 3.35, p = .002, BF_10_ = 16.00, Figure 3B) and errors (t(28) = 3.00, p = .006, BF_10_ = 6.77, Figure 3C).

**Figure 3.**
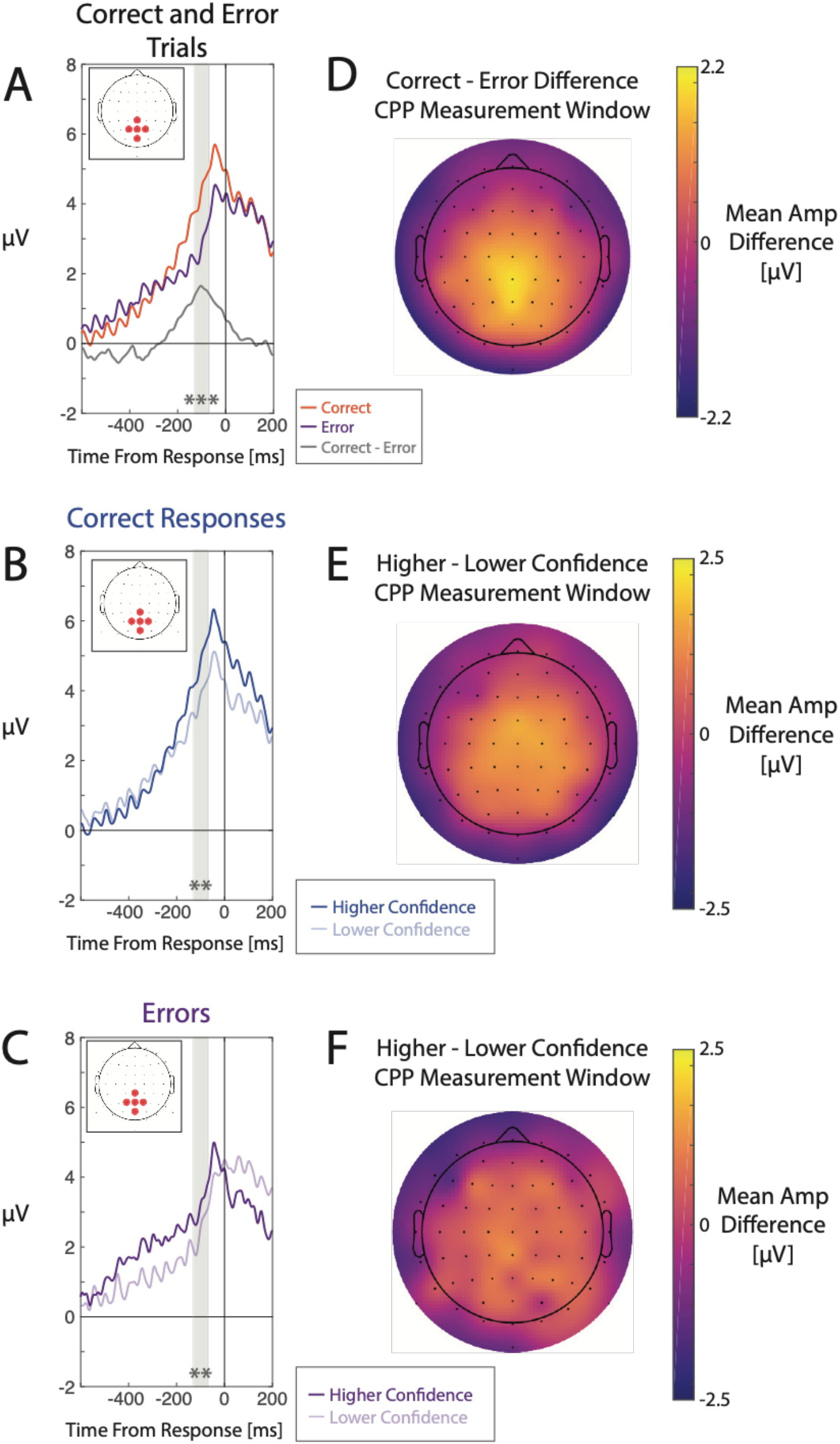
ERPs time-locked to the response to the S1 target at channels Pz, P1, P2, CPz, and POz. A) ERPs for correct and error responses at the parietal region of interest (ROI). Grey shaded regions denote the mean amplitude measurement time windows for the CPP. B) ERPs for higher/lower confidence ratings in trials with correct responses. C) ERPs for higher/lower confidence ratings in trials with errors. D, E, F) Scalp maps of mean amplitudes across the CPP time window, for each of the across condition contrasts displayed in A-C. All ERPs were baseline-corrected using a pre-stimulus baseline. Asterisks denote statistically significant differences between correct responses and errors or significant associations between confidence ratings and ERP component amplitudes (** denotes p < .01 and *** denotes p < .001).

We additionally plotted heat maps of single trial amplitudes sorted by RT to verify whether the CPP was actually response-locked in our data (as done by O’Connell et al., 2012; van Vugt et al., 2019). These heat maps revealed a positive-going component (i.e., the CPP) that was closely aligned to the time of the response (plotted in Supplementary Figure S5).

To aid comparison with existing work (e.g., Gherman & Philiastides, 2018; Rausch et al., 2020), we have plotted stimulus-locked ERPs for correct/error trials and higher/lower confidence ratings in Supplementary Figure S6. We also analysed stimulus-locked CPP mean amplitude measures. In these analyses, we found that CPP amplitudes were larger for correct compared to erroneous responses (t(28) = 8.33, p < .001, BF_10_ > 2.71 * 10^6^, Figure S6A). CPP amplitudes were associated with confidence for trials with correct responses (t(28) = 4.59, p < .001, BF_10_ > 290, Figure S6B) but not for trials with errors (t(28) = -1.58, p = .125, BF_10_ = 0.60, Figure S6C). Based on our assumption that the CPP reflects a response-locked ERP component, we have not focused on these results in our paper. However, we acknowledge that they might be interesting to other researchers who assume that the CPP is better described as a stimulus-locked component.

#### 3.2.1. Effects of Current Source Density Transformation

After observing positive associations between CPP amplitudes and confidence we repeated these analyses using CSD-transformed data. This approach follows Kelly and O’Connell (2013) who observed similar effects of stimulus discriminability (which closely covaries with confidence) on CPP pre-response amplitudes. When they applied a CSD transformation, there was little evidence of an association between CPP amplitudes and decision confidence. Response-locked CSD-ERPs are plotted in Figure 4. CPP amplitudes remained larger for correct responses as compared to errors (t(28) = 5.29, p < .001, BF_10_ > 1,700, Figure 4A, left panel). Associations between CPP amplitudes and confidence were no longer observed for trials with correct responses (t(28) = 0.45, p = .656, BF_10_ = 0.22, Figure 4B, left panel) or trials with errors (t(28) = 1.61, p = .118, BF_10_ = 0.63, Figure 4C, left panel).

**Figure 4.**
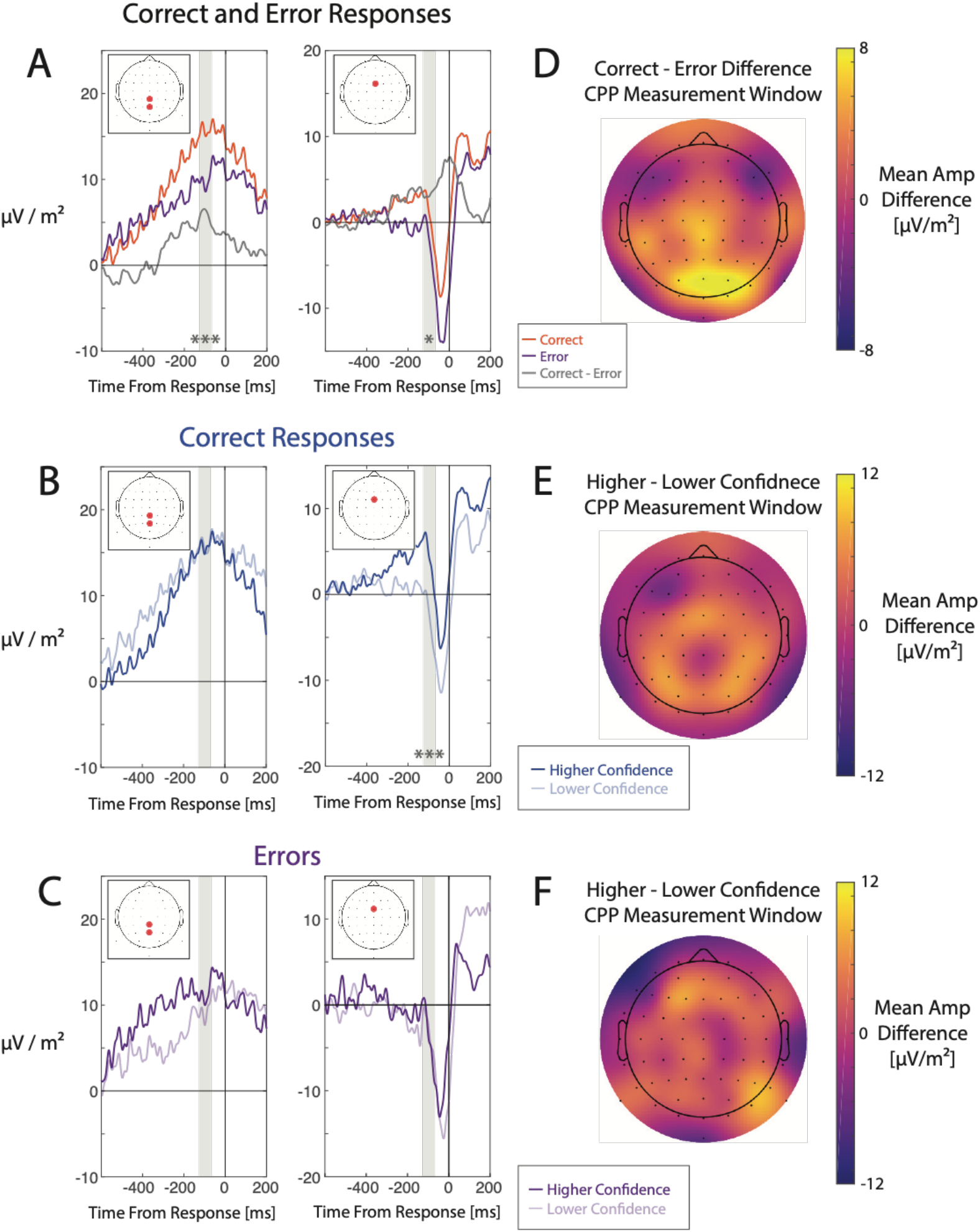
CSD-transformed ERPs time-locked to the response to the S1 target at parietal channels Pz and CPz (left ERP plot panels) and fronto-central channel FCz (right ERP plot panels). A) CSD-ERPs for correct and error responses. Grey shaded regions denote the mean amplitude measurement time windows used for measuring the CPP and frontal component. B) ERPs for higher/lower confidence ratings in trials with correct responses. C) ERPs for higher/lower confidence ratings in trials with errors. D, E, F) Scalp maps of mean amplitudes across the CPP time window, for each of the across-condition contrasts displayed in parts A-C. All ERPs were baseline-corrected using a pre-stimulus baseline. Asterisks denote statistically significant differences between correct responses and errors or significant associations between confidence ratings and ERP component amplitudes (* denotes p < .05 and *** denotes p < .001).

We also repeated our CPP analyses using the parietal ROI electrodes that were included in the non CSD-transformed ERP analyses. The results aligned well with the analyses that included only Pz and CPz. Mean amplitudes were larger for correct responses compared to errors (t(28) = 7.13, p < .001, BF_10_ > 160,000). We did not find statistically significant associations between CPP amplitudes and confidence for trials with correct responses (t(28) = 1.75, p = .090, BF_10_ = 0.76) or errors (t(28) = 1.85, p = .074, BF_10_ = 0.89).

To assess whether the apparent effects on the CPP (in the non CSD-transformed data) were actually due to amplitude modulations of a frontal component (as reported in Kelly & O’Connell, 2013) that spread to parietal electrodes via volume conduction, we additionally measured mean amplitudes at electrode FCz using the CPP measurement time window. Heat maps of single trial amplitudes sorted by RT revealed that this frontal component was closely time-locked to the response in our data (see Supplementary Figure S5). Mean amplitudes were larger for correct responses compared to errors (t(28) = 2.45, p = .021, BF_10_ = 2.46, Figure 4A, right panel). Positive associations with confidence were observed for trials with correct responses (t(28) = 5.11, p = < .001, BF_10_ > 1,100, Figure 4B, right panel) but not for errors (t(28) = 1.30, p = .203, BF_10_ = 0.42, Figure 4C, right panel). Notably, these effects in correct response trials appeared to occur from ∼250 ms prior to the response.

To further investigate the influence of the frontal component on non CSD-transformed ERP measures at parietal electrodes, we performed regression analyses using confidence ratings to predict mean amplitudes during the CPP pre-response time window for each electrode separately (similar to Kelly & O’Connell, 2013). Scalp maps of group-averaged beta values, intercepts and mean amplitudes are plotted in Supplementary Figure S7. For trials with correct responses, the topography of beta values (showing where associations with confidence were strongest) was focused over Cz and FCz, and was more anterior than the distribution of intercepts and mean amplitude values, which resembled the typical parietal topography of the CPP component (e.g., Twomey et al., 2015).

Stimulus-locked ERPs are also plotted in Supplementary Figure S8. Stimulus-locked CPP amplitudes were larger for correct compared to erroneous responses (t(28) = 5.27, p < .001, BF_10_ > 1,600, Figure S8A). We did not find associations between confidence and CPP amplitudes for trials with correct responses (t(28) = 1.33, p = .195, BF_10_ = 0.44, Figure S8B) or errors (t(28) = 0.53, p = .603, BF_10_ = 0.22, Figure S8C).

### 3.3. Pe Component Amplitudes

We analysed Pe component mean amplitudes (between 200-350 ms from the time of the response) using both pre-stimulus and pre-response ERP baselines in separate analyses. This was done to systematically assess whether the use of pre-response baselines artificially produces observed associations between Pe amplitudes and confidence in cases where there are already ERP differences across conditions prior to the response (e.g., in Boldt & Yeung, 2015). Both sets of analyses only included trials in which the informative S2 stimulus did not appear.

#### 3.3.1. Analyses Using Pre-Stimulus Baseline-Corrected ERPs

We first measured mean amplitudes of the Pe component using a pre-stimulus baseline, which are not influenced by ERP differences across conditions that might already exist prior to the response (as we discuss in detail in section 1.2). Using this type of baseline correction, we did not observe Pe amplitude differences between trials with correct responses and trials with errors (t(28) = 0.98, p = .335, BF_10_ = 0.31, Figure 5A). In contrast to the CPP results, for trials with correct responses, mean Pe amplitudes over this time window were not associated with decision confidence (t(28) = 0.45, p = .656, BF_10_ = 0.22, Figure 5B). Bayes factors of < 0.3 indicated a moderate amount of evidence for the null hypothesis, meaning that we did not replicate the Pe-confidence association for correct response trials reported by Boldt and Yeung (2015). For trials with errors, Pe amplitudes were negatively associated with decision confidence, with larger (i.e. more positive) Pe amplitudes observed in trials with lower confidence (t(28) = 2.82, p = .009, BF_10_ = 5.12, Figure 5C). The observed association for errors, but not for trials with correct responses, is consistent with patterns of pre-stimulus baseline-corrected Pe amplitudes plotted in Boldt and Yeung (2015, their Figure 3B).

**Figure 5.**
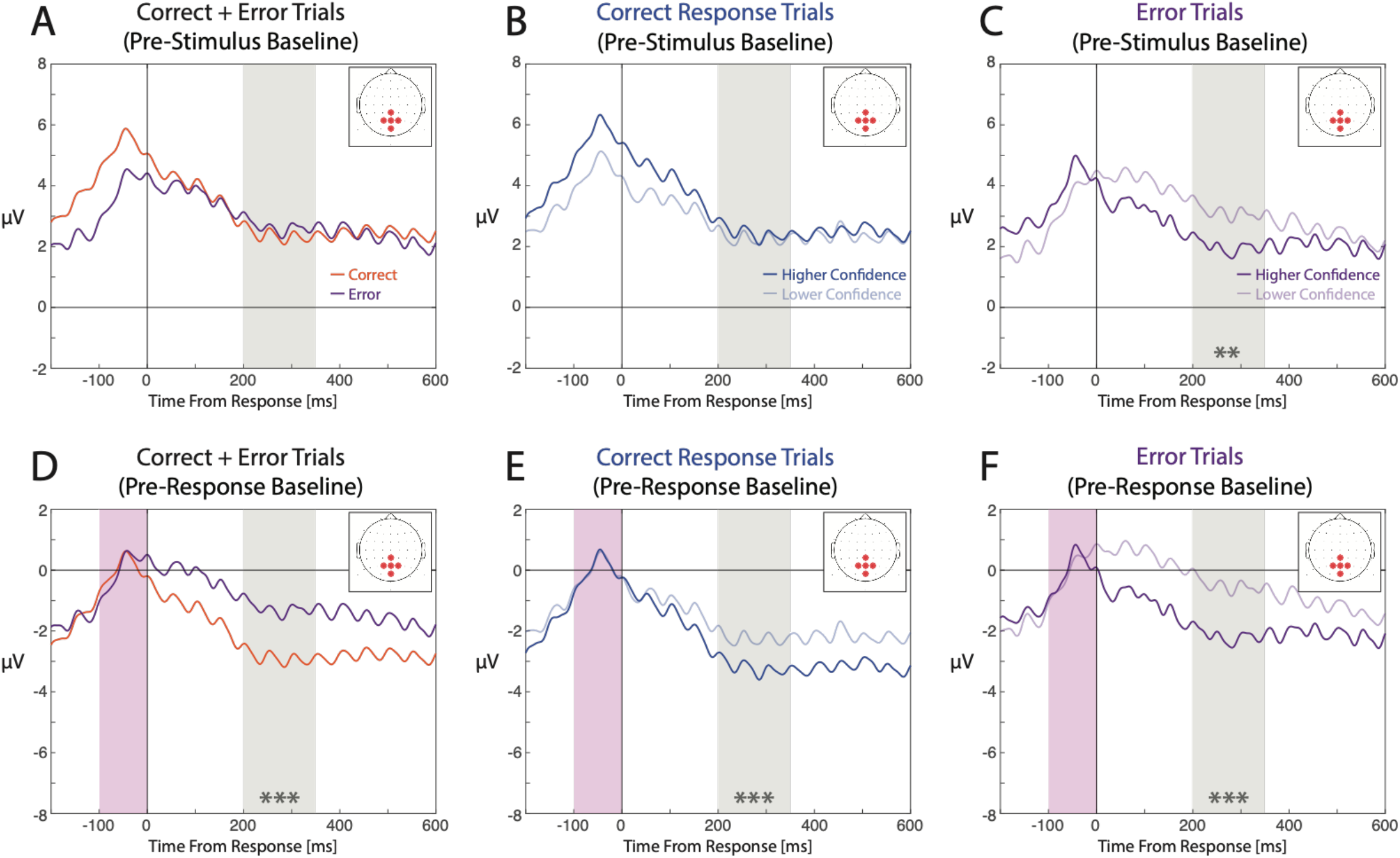
ERPs following responses to S1 targets at parietal ROI electrodes Pz, P1, P2, CPz, and POz. ERPs corrected using a pre-stimulus baseline are shown in A-C (top row). ERPs corrected using a pre-response baseline are shown in D-F (bottom row). A, D) ERPs for trials with correct and erroneous responses. B, E) ERPs for higher/lower confidence ratings in trials with correct responses. C, F) ERPs for higher/lower confidence ratings in trials with errors. In all plots the grey shaded area denotes the 200-350 ms time window used to measure the Pe component. The shaded magenta area denotes the pre-response baseline time window. Asterisks denote statistically significant differences between correct responses and errors or significant associations between confidence ratings and ERP component amplitudes (** denotes p < .01 and *** denotes p < .001).

#### 3.3.2. Analyses Using Pre-Response Baseline-Corrected ERPs

We also repeated our analyses using a pre-response baseline (−100 to 0 ms relative to the response) to compare our results to those of Boldt and Yeung (2015). In contrast to analyses of pre-stimulus baseline-corrected ERPs, Pe amplitudes for trials with errors were more positive-going compared to those with correct responses (t(28) = -5.21, p < .001, BF_10_ > 1,300, Figure 5D). There were clear negative-going associations between Pe amplitudes and confidence ratings both for trials with correct responses (t(28) = -4.48, p < .001, BF_10_ > 230, Figure 5E) and trials with errors (t(28) = -4.83, p < .001, BF_10_ > 550, Figure 5F). Notably, the apparent timing and duration of these effects are almost identical to the ERP differences across confidence rating categories in Boldt and Yeung (2015, their Figure 3A).

Taken together, the results from analyses of pre-stimulus and pre-response baseline-corrected ERPs show that, when there are already ERP differences across conditions prior to the response (as indicated by effects on the CPP), the use of a pre-response baseline produces artefactual associations between Pe amplitudes and confidence. Using the more appropriate pre-stimulus baseline correction, however, these results demonstrate that Pe amplitudes were only truly associated with variations in confidence in error trials.

### 3.4. Analyses Using Restricted Ranges of Confidence Ratings

We also performed complimentary, post hoc regression analyses using restricted ranges of confidence ratings, including the range from “unsure” (0) to “certainly correct” (100; indexing participants’ certainty that they had made a correct response) and, in separate analyses, the range from “certainly wrong” (−100) to “unsure” (indexing participants’ certainty that they had made an error). Results are detailed in the Supplementary Material.

In summary, the results were broadly consistent with those of the main analyses. pre-response amplitudes were positively associated with confidence across the range of “unsure” to “certainly correct” for the CPP in conventional (i.e., non CSD-transformed) ERPs, but not for CSD-transformed ERPs. Frontal Component amplitudes were positively associated with confidence across the range of “unsure” to “certainly correct”. For confidence ratings across the range of “certainly wrong” to “unsure”, a negative-going association was found, however the Bayes factor of BF_10_ = 1.38 indicated only weak evidence in favour of the alternative hypothesis. Pe amplitudes were associated with confidence across the range of “certainly wrong” to “unsure” when using both pre-stimulus and pre-response baselines. However, associations with confidence across the range of “unsure” to “certainly correct” were only found for the Pe when using pre-response baselines.

## 4. Discussion

To characterise the electrophysiological correlates of confidence we presented participants with a challenging perceptual decision task. We varied stimulus discriminability (i.e., target contrast) over a wide range, which produced marked variation in self-reported levels of confidence. We identified ERP correlates of confidence both during decision formation and after a decision had been made. By analysing conventional (non CSD-transformed) ERPs, we found that the amplitude of the response-locked CPP component positively correlated with decision confidence. Subsequent analyses using CSD-transformed data, however, did not find evidence for this association, and instead provided moderate evidence in favour of the null hypothesis (BF_10_= 0.22). Analyses of activity at electrode FCz revealed that associations observed in the non CSD-transformed data may instead be attributable to a frontal ERP component that influenced measures at parietal electrodes via volume conduction.

We also tested for associations between confidence and amplitudes of the post-decisional Pe component using a pre-stimulus baseline, and ran the same analyses using the conventional pre-response ERP baseline. Importantly, effects on ERPs corrected using a pre-response baseline were likely to reflect signals of a pre-response origin rather than a true modulation of the Pe component. Indeed, when we used a pre-response baseline, there was a strong negative association between confidence and Pe amplitudes for both trials with correct responses and errors. However, when using a more appropriate pre-stimulus baseline, we found this association for trials with errors, but not for trials with correct responses. In the latter case, the Bayes factor provided moderate evidence in favour of the null hypothesis (BF_10_= 0.22).

Our findings, which are not subject to the same methodological issues as previous work, encourage a re-evaluation of existing evidence that links ERPs and decision confidence. They suggest that certainty in having made a correct decision is indexed by fronto-central activity during the evidence accumulation stage, whereas certainty in having committed an error is indexed by the amplitude of the post-decisional Pe component. By extension, it appears that confidence does not correlate with any single ERP component in a consistent direction across the rating spectrum ranging from ‘certainly wrong’ to ‘unsure’ to ‘certainly correct’. Instead, confidence judgments may be jointly informed by processes occurring over distinct pre-and post-decisional time windows, with one’s degree of confidence in favour of a correct decision being computed during decision formation, and error detection occurring after a decision has been made (as proposed by Rausch et al., 2020). Importantly, our findings are not fully compatible with existing theoretical accounts that use ERP findings to claim preferential support for decisional locus and post-decisional locus models of confidence (e.g., Philiastides et al., 2014; Boldt & Yeung, 2015; Desender et al., 2021b).

### 4.1. Neural Correlates of Confidence During Decision Formation

Our analyses of conventional (i.e., non CSD-transformed) ERPs revealed that CPP amplitudes (measured at centro-parietal channels) were positively associated with confidence ratings for both trials with correct responses and errors. This pattern, seen in our response-locked ERPs, aligns with existing studies that have measured stimulus-locked CPP/P3 amplitudes (e.g., Squires et al., 1973; Gherman & Philiastides, 2015, 2018; Zarkewski et al., 2019; von Lautz et al., 2019; Herding et al., 2019; Rausch et al., 2020), as well as Philiastides et al. (2014) who found associations between response-locked CPP amplitudes and model-derived (rather than self-reported) confidence ratings. By analysing CPP amplitudes time-locked to the behavioural response, we verified that the association between CPP amplitudes and self-reported confidence was not simply due to differences in the timing of the CPP component across confidence rating conditions (also see section 1.1 above). Our findings demonstrate the utility of including both stimulus- and response-locked ERP measures that provide complimentary information about an ERP component.

Based on the observed accumulation-to-threshold morphology of the CPP, researchers have interpreted larger CPP amplitudes as reflecting a greater degree of accumulated evidence in favour of the chosen option in trials with higher confidence ratings (e.g., Philiastides et al., 2014; Gherman & Philiastides, 2015; von Lautz et al., 2019). This has been taken as support for the ‘balance-of-evidence hypothesis’ as specified in some decisional locus models (e.g., Vickers, 1979; Vickers & Packer, 1982; Ratcliff & Starns, 2009), which specifies that confidence indexes differences in the positions of racing accumulators in discrete choice tasks. However, when we applied a CSD-transform to our data we no longer found associations between CPP amplitudes and confidence, with the Bayes factor for analyses of trials with correct responses (BF_10_ = 0.22) indicating moderate evidence in favour of the null hypothesis. Instead, we found that the CPP-confidence associations identified using non CSD-transformed ERPs may have been due to a temporally overlapping frontal component whose amplitude positively correlated with confidence. Importantly, this frontal component appeared to bias ERP amplitude measurements at centro-parietal channels via volume conduction (as found in Kelly & O’Connell, 2013; see also Dmochowski & Norcia, 2015). Our findings therefore cast doubt on the idea that CPP/P3 amplitudes are a genuine correlate of decision confidence. Despite the CPP previously being closely linked to evidence accumulation dynamics (Twomey et al., 2015; Kelly et al., 2021), our findings suggest that the amplitude of the frontal component may instead reflect the extent of evidence accumulated in favour of the selected choice option as specified in decisional locus models.

Here, we note that our findings are broadly congruent with decisional-locus models (which were developed using behavioural data and do not specify ERP correlates of evidence accumulation). The fact that confidence for correct responses, but not errors, was correlated with frontal component amplitudes is also consistent with decisional-locus models. If participants thought they were committing an error at the time of the decision in our difficult perceptual discrimination task, they would presumably have changed their decision. Therefore, it is reasonable that error detection (indexed by the Pe) would occur only after the response had been made.

Although frontal component amplitudes positively correlated with confidence ratings, it is unclear whether this reflects processes that are specifically associated with confidence computations. For example, Kelly and O’Connell (2013) found that amplitudes of this component correlated with RT, which often covaries with confidence in perceptual decision tasks (e.g., Johnson, 1939; Festinger, 1943; Vickers & Packer, 1982; Kiani et al., 2014). Kelly and O’Connell (2013) likened this component to movement preparation-related components such as the Contingent Negative Variation or the Bereitschaftspotential (Brunia and van Boxtel, 2001; Baker et al., 2012). Given that some aspects of motor action execution (such as response force) covary with confidence (e.g., Gajdos et al., 2019; reviewed in Turner et al., 2021), it is not surprising that ERP components linked to motor action execution might also correlate with confidence. To better understand the relationships between fronto-central activity, response speed and confidence, it may be useful to investigate graded variations in confidence that are not closely correlated with RT (e.g., using similar designs to Bang & Fleming 2018; Fleming et al., 2018) or use model-based approaches that explicitly account for differences in confidence across fast and slow RTs (e.g., Rausch et al., 2020).

We also caution that we may not have had sufficient statistical power to detect associations between confidence and frontal component amplitudes in trials with errors, as there were smaller numbers of these trials compared to correct responses. Our post-hoc analyses (that included both correct and error trials) identified positive-going associations for confidence ratings within the range of “unsure” to “certainly correct”, but weak evidence (BF_10_ = 1.38) for a negative-going association across the range of “certainly wrong” to “unsure”. This suggests that frontal component amplitudes may actually scale with certainty in having made a correct response or an error, rather than confidence per se (for a distinction between these concepts see Pouget et al., 2016). Future work should investigate the relationship between frontal component amplitudes and confidence ratings indicating that an error had occurred, to better characterise any possible links between this ERP component, certainty, and error detection.

We additionally note that, in the CSD-ERPs, there were apparent differences between higher/lower confidence ratings prior to the CPP measurement window (Figures 4B, 4C, left panels). These ERP effects reflect differences in the CPP build-up rate across subsets of trials with faster and slower RTs, as typically observed in similar perceptual decision tasks (e.g., O’Connell et al., 2012; Kelly & O’Connell, 2013; Twomey et al., 2015; Feuerriegel et al., 2021a). The build-up rate of the CPP is thought to index the *rate* of evidence accumulation, known as the drift rate in evidence accumulation models (O’Connell et al., 2012; Kelly et al., 2021). By contrast, CPP pre-response amplitudes are interpreted here as the *extent* of evidence accumulation at the time of decision commitment. It is therefore important to choose amplitude measurement windows that are not largely affected by differences in CPP build-up rates. We selected our time window of -130 to -70 ms relative to the response based on previous work that identified this time window as suitable for this purpose (Steinemann et al., 2018; Kelly et al., 2021; Feuerriegel et al., 2021). Please also note that, although CPP build-up rates are often important to consider in decision-making research (e.g., O’Connell et al., 2012; Kelly et al., 2021), we did not measure CPP slopes as they were not relevant to claims about the *extent* of evidence accumulation as specified in decisional-locus models.

### 4.2. Post-Decisional Correlates of Confidence

We systematically tested for associations between Pe amplitudes and confidence using pre-stimulus and pre-response baselines in separate analyses. We analysed non CSD-transformed ERPs to be consistent with prior work on the Pe component and decision confidence. We found that, when using a pre-stimulus baseline, Pe amplitudes (measured at centro-parietal electrodes) inversely scaled with confidence in trials with decision errors, but not in trials with correct responses. In the latter case, the Bayes factor (BF_10_ = 0.22) indicated moderate evidence in favour of the null hypothesis. However, when we used a pre-response baseline, we replicated previous reports of more positive-going Pe amplitudes in trials with lower confidence ratings, for both correct responses and errors (Boldt & Yeung, 2015). This difference in patterns of results is because amplitudes at the same centro-parietal electrodes were already positively correlated with confidence during the pre-response period (indexed by effects on CPP component amplitudes).

These findings demonstrate that ERP differences which occur before the response can be mistakenly interpreted as amplitude differences of post-response ERP components (such as the Pe) when pre-response baselines are used. The reason for this is that a pre-response baseline correction will nullify existing differences in the respective baseline time window and - if these differences are systematically related to the conditions of interest - artificially propagate them into subsequent time windows. This suggests that associations between confidence and Pe amplitudes (in correct response trials) reported in Boldt and Yeung (2015, see also Desender et al., 2019b) may reflect differences in pre-response CPP amplitudes across confidence ratings, rather than true differences in Pe amplitudes. However, our results are broadly congruent with the pattern of Pe amplitudes visible when using a pre-stimulus baseline in Boldt and Yeung (2015), whereby Pe amplitudes increased with higher certainty in having made an error, but not with higher certainty in having made a correct response (their Figure 3B, see also Desender et al., 2019b).

These findings run contrary to a recently proposed model that attempts to unify error detection and decision confidence into a single framework (Desender et al., 2021b). According to this model, two-choice decisions are initially made according to a double-bounded evidence accumulation process. Following the decision, the ‘reference frame’ of an ensuing metacognitive decision is proposed to shift to a single-bounded accumulation process that reflects one’s degree of evidence that a decision error has been committed. In other words, the decision is framed similarly to a single-bounded stimulus detection decision (e.g., O’Connell et al., 2012), where a decision error is the event to be detected. Based on the findings of approximately monotonic relationships between decision confidence and Pe amplitudes in Boldt and Yeung (2015), this model proposes that one’s degree of decision confidence is computed based on the extent of post-decisional evidence that has been accumulated in favour of making an error. Importantly, Desender et al. (2021b) claim that the extent of accumulated post-decisional evidence is reflected in the amplitude of the Pe component, and that the amplitude of the Pe component should show a monotonic, inverse relationship with decision confidence ratings (see their Figure 1C). By framing post-decisional evidence accumulation in this way, the model fits error detection and confidence judgments (ranging from ‘certainly incorrect’ to ‘unsure’ to ‘certainly correct’) into the same underlying framework.

Contrary to the assumptions of the model, we did not find evidence supporting the notion that decision confidence shows a simple monotonic relationship with Pe amplitudes across the full confidence rating spectrum. Rather, it appears that Pe amplitudes scale with one’s degree of certainty that they had made an error, specifically in trials where an error had been committed. Importantly, we did not observe evidence of covariation between Pe amplitudes and confidence ratings for trials with correct responses. This pattern more closely resembles a hypothetical evidence accumulation associated with error detection (e.g., Murphy et al., 2015) rather than decision confidence more generally. This in turn suggests that decision confidence and error detection do not neatly fit into the single framework proposed by Desender et al. (2021b).

Our findings hint at dissociable sources of information being used to compute confidence for correct responses and for errors. However, we caution that it is unclear whether these effects on ERP components reflect computations that are critical to our sense of confidence, or changes to other decision processes that co-vary with confidence ratings. For example, errors typically constitute a rare and surprising event when performance is well above chance (Wessel & Aron, 2017). When errors are detected, this triggers a cascade of processes that onset rapidly after the error, for example those associated with the orienting response (reviewed in Wessel, 2018).

Consequently, it is unclear whether Pe amplitudes in our study (and other paradigms with similar properties) reflect different proportions of detected errors (and associated surprise-related responses) across confidence rating conditions. Formal models of error detection and confidence (e.g., Desender et al., 2021a) may be useful for identifying patterns that are more specifically related to decision confidence.

We additionally note that the baseline-related issue described above is not particular to Boldt and Yeung (2015) and is present in the work of many others who have investigated post-decisional ERP correlates of error detection and confidence (e.g., Selimbelogyu et al., 2012; Desender et al., 2019a, 2019b; Rausch et al., 2020).

### 4.3. Study Limitations

Our findings should be interpreted with the following caveats in mind. Firstly, our experiment design is different to previous work in that, in 50% of trials, an S2 target (which was informative regarding the correct response in the trial) appeared after responding to the S1 target. Although the appearance of this stimulus was not predictable, it is possible that participants anticipated the onset of the S2 stimulus, which could help them make more accurate confidence ratings in trials where they were unsure of their decision (i.e., had lower confidence). Based on this idea, it could be argued that the frontal component identified in our study reflects the focusing of attention in preparation for the S2 stimulus rather than confidence. We do not believe this is the case because a similar frontal component was observed in Kelly and O’Connell (2013), and they did not present informative S2 stimuli. However, we recommend that future work attempts to identify this frontal component in situations where there is no anticipation of upcoming task-relevant information (or even task performance feedback).

We also note that, based on analyses of our own data, we cannot be certain that the same patterns of results will be found in re-analyses of existing datasets. For example, the study of Boldt & Yeung (2015) did not use post-target masking, and sensory information may have been available for post-decisional evidence accumulation to a greater extent than in our experiment, or others that used post-target masks (e.g., Rausch et al., 2020; for discussion of the dynamics of post-decisional evidence accumulation see Resulaj et al., 2009; Turner et al., 2022). In addition, we used interleaved, continuously-running staircases to determine target contrast, which differs to previous work that used a single stimulus discriminability level (e.g., Boldt & Yeung, 2015) or multiple, discrete levels (e.g., Rausch et al., 2020). Our inferences here are based on the fact that we replicated effects seen in previous work when using similar analysis methods to those studies, but found markedly different results when using other analysis methods that avoid the issues mentioned above. For example, the lack of Pe amplitude variation across confidence ratings in favour of a correct decision mirrors the apparent lack of Pe amplitude differences when using a pre-stimulus baseline in Boldt & Yeung (2015, their Figure 3B). However, we believe that our analysis approach should be applied to a range of existing datasets to assess whether our results generalise across different stimulation and task contexts, as well as different confidence rating scales.

In addition, we found that participants varied in how they used the confidence rating scale. Although mean confidence ratings positively scaled with stimulus discriminability (depicted in Figure 2B) and group-averaged confidence ratings showed similar patterns to previous work (e.g., Fleming et al., 2018; Turner et al., 2021), some participants provided a much broader range of confidence ratings than others (shown in Supplementary Figure S3). For our dataset (and many others), it is difficult to know whether inter-individual differences in confidence rating distributions reflect actual differences in internal estimates of confidence, or differences in how such internal estimates map onto the ratings given by participants (known as the criterion problem, see Peters & Lau, 2015). Because of this, we were restricted to testing for linear relationships between confidence and ERP component amplitudes (following the analysis approach of Boldt & Yeung, 2015). Consequently, we may have missed more complex, non-linear relationships between ERP amplitudes and confidence ratings across the scale (ranging from “certainly wrong” to “unsure” to “certainly correct”) that may be associated with certainty rather than confidence (for a distinction between these concepts see Pouget, 2016). Future work seeking to identify fine-grained non-linear relationships between confidence and neural measures could employ strategies that promote a standardised use of the entire confidence rating scale, although this may require extensive training prior to the experiment.

We also note that the frontal component identified in our study (which had an onset of ∼250 ms prior to the response) appeared to overlap in time with the later error-related negativity (ERN) component (Falkenstein et al., 1991; Gehring et al., 1993). The ERN is typically more negative-going following commission of a decision error as compared to a correct response (Gehring et al., 1993; Bode & Stahl, 2014; but see Di Gregorio et al., 2018), and has been investigated as a possible neural correlate of confidence (e.g., Boldt & Yeung, 2015; Rausch et al., 2020). Although we measured the frontal component over a time window earlier than that used to measure the ERN (e.g., -40 to 60 ms relative to the response in Boldt & Yeung, 2015), we could not accurately measure the ERN itself due to the overlap. Further work is needed to clearly define the extent of covariance between these two components, in order to ascertain whether they reflect similar processes during the time-course of decision formation. If they do reflect distinct sources of neuronal activity, then measuring the ERN using a pre-response baseline window that overlaps with the frontal component (e.g., as in Boldt & Yeung, 2015; Selimbelogyu et al., 2012; Desender et al., 2019a, 2019b; Rausch et al., 2020) may influence ERN amplitude measures. This may be problematic if activity at fronto-central electrodes also differs across conditions of interest prior to the response (e.g., Kelly & O’Connell, 2013, see Figure 4 above).

There are also two factors to consider when comparing our non CSD-transformed and CSD-transformed results. The first is that CSD-transformation attenuates sources of neural activity that are broadly-distributed across the scalp (Kayser & Tenke, 2006). The CPP is reliably found in CSD-transformed data and shows accumulation-to-bound trajectories that are characteristic of this ERP component (Kelly & O’Connell, 2013; Steinemann et al., 2018; Feuerriegel et al., 2021a; Kelly et al., 2021). However, we cannot rule out the possibility that there may have been more broadly-distributed sources of ERPs that covary with confidence and were attenuated by CSD transformation. Whether these (if they exist) can be classified as the CPP component, however, is unclear. In any case, the topographies of associations between confidence and ERP amplitudes during the pre-response CPP time window in Supplementary Figure S6 show that the frontal component identified in our study is very likely to bias measures in non CSD-transformed data, and caution should be taken to dissociate any overlapping effects.

The second factor is that CSD-transformed ERP measures tend to be more variable compared to non CSD-transformed ERPs (Vidal et al., 2003). This may have prevented us from identifying associations between CPP amplitudes and confidence in CSD-transformed data. However, we believe this is unlikely, as beta coefficients were tightly clustered around zero (Supplementary Figure S4), and the Bayes factor of BF_10_ = 0.22 indicated moderate evidence for the null hypothesis, rather than showing values around 1 that do not provide substantial support for the null or alternative hypotheses, as would be expected if only the variability had increased. However, we note that this Bayes factor does not indicate overwhelming evidence for the null, and analyses of existing datasets will be useful to see if this null result can be replicated.

Finally, our analyses of response-locked CPP component amplitudes rely on the assumption that the CPP is in fact closely time-locked to the keypress response, in line with the original definition of the CPP (O’Connell et al., 2012). Notably, others have considered this component as stimulus-locked (e.g., Rausch et al., 2020). We encourage researchers to verify whether the component is in fact time-locked to the response in their own datasets. We also note that combining stimulus- and response-locked analyses of CPP amplitudes may provide useful complimentary information about how this component covaries with factors such as decision confidence.

### 4.4. Conclusion

We probed the neural correlates of decision confidence using a difficult perceptual discrimination task. By analysing conventional (non CSD-transformed) ERPs we confirmed that pre-response CPP amplitudes are correlated with confidence. However, analyses of CSD-transformed ERPs revealed that these effects at centro-parietal channels might be due to the influence of a frontal component whose amplitude was also correlated with confidence. This frontal effect appeared to influence measures of the CPP at centro-parietal channels via volume conduction. By systematically analysing the post-decisional Pe component using pre-stimulus and pre-response baselines, we also determined that the amplitude of the Pe inversely scaled with confidence, but we only observed this association in trials with erroneous decisions. Our findings highlight the possibility that previously reported relationships between Pe component amplitudes and the full spectrum of confidence across correct and error trials were (at least partly) due to methodological issues related to the use of pre-response baselines. Taken together, our findings suggest that certainty in having made a correct decision is indexed by fronto-central activity during decision formation, and certainty in having made an error is indexed by the amplitude of the post-decisional Pe component. These processes, which occur over distinct time windows, may jointly inform confidence judgments in perceptual decision tasks.

## Acknowledgements

This project was supported by an Australian Research Council Grant (ARC DP160103353) to S.B. and R.H and an Australian Research Council Discovery Early Career Researcher Award to D.F. (ARC DE220101508). Funding sources had no role in study design, data collection, analysis or interpretation of results. We thank Pragya Arora for her help with data collection.

## Notes

https://osf.io/gazx2/

### Competing Interest Statement

The authors have declared no competing interest.

