## Supplementary Material for "Electrophysiological correlates of confidence differ across correct and erroneous perceptual decisions"

### Mixed Effects Regression Model Structures and Coefficients for Analyses of Accuracy, Response Times, and Confidence Ratings

Code for fitting all models listed below will be available at https://osf.io/gazx2/ at the time of publication. To test for fixed effects of each variable of interest on accuracy, response time (RT) and confidence rating measures, we compared models with and without the fixed effect of interest, but with identical random effects structures, using likelihood ratio tests. Models were fit using the R package lme4 (Bates et al., 2015). We used a random effects structure that included a random intercept by participant and random slopes of S1 target contrast (for analyses of accuracy and RTs) and trial outcome (correct/error, for analyses of RTs).

#### List of Variable Names and Definitions

`RT` – The response time following S1 targets in seconds

`trial_outcome` – The decision outcome following S1 targets (correct or error)

`confidence_normalised` – Decision confidence ratings, normalised to be between 0-1 (with 0 corresponding to certainly wrong, i.e., the minimum confidence rating, and 1 corresponding to certainly correct, i.e., the maximum confidence rating).

`target_contrast` – The contrast level of the dominant grating in the S1 target (ranging between 0.5 and 1). This is also referred to as target discriminability in the corresponding paper.

`ID` – The participant ID number (used for defining random intercepts and slopes)

#### Regression Model Equations and Coefficients

Full model equations are listed below. Here, we follow the conventions of the lme4 package notation (Bates et al., 2015) when describing the structure of mixed effects regression models. Model coefficients for the full model (including the fixed effect of interest) are displayed below each set of model equations.

#### Testing for an Effect of S1 Target Contrast (i.e., Discriminability) on RTs

For analyses of S1 RTs, we fit generalised linear mixed effects regression models (Gamma family) with an identity link function. We included both trial where the S2 target did and did not appear, as the presentation of the S2 target occurred after the response to the S1 target had been made. We compared the following model including effects of S1 target contrast:

```
RT ~ target_contrast + trial_outcome + (1 + target_contrast +
trial_outcome | ID)
```

With a comparison model that did not include a fixed effect of S1 target contrast:

```
RT ~ trial_outcome + (1 + target_contrast + trial_outcome | ID)
```

Coefficients for the generalised linear mixed model (Gamma family, identity link function) fit by maximum likelihood

| | | | | | |
|---|---|---|---|---|---|
| | AIC | BIC | Log Likelihood | Deviance | |
| | -13457.0 | -13382.3 | 6738.5 | -13477.0 | |
| Scaled Residuals: | | | | | |
| | Min | 1Q | Median | 3Q | Max |
| | -2.812 | -0.634 | -0.099 | 0.560 | 5.620 |
| Random Effects: | | | | | |
| | Groups | Name | Variance | Std. Dev | |
| | ID | (Intercept) | 0.011 | 0.103 | |
| | | target_contrast | 0.017 | 0.129 | |
| | | trial_outcome | 0.001 | 0.030 | |
| | Residual | | 0.085 | 0.291 | |
| Number of observations: 12920, groups: ID, 29 | | | | | |
| Fixed Effects | | | | | |
| | | Estimate | Std. Error | t value | |
| | (Intercept) | 0.834 | 0.042 | 19.79 | |
| | target_contrast | -0.377 | 0.055 | -6.84 | |
| | trial_outcome | -0.037 | 0.012 | -3.07 | |

**Testing for an Effect of S1 Target Contrast (i.e., Discriminability) on Proportions of Correct Responses to S1 Targets**

For analyses of proportion correct responses (i.e. accuracy) for S1 targets, we fit generalised linear mixed effects regression models (Binomial family) with a logit link function. We included both trial where the S2 target did and did not appear, as the presentation of the S2 target occurred after the response to the S1 target had been made. We compared the following model including effects of S1 target contrast:

```
trial_outcome ~ target_contrast + (1 + target_contrast | ID)
```

With a comparison model that did not include a fixed effect of congruency:

```
trial_outcome ~ (1 + target_contrast | ID)
```

Coefficients for the generalised linear mixed model (Binomial family, logit link function) fit by maximum likelihood

| | | | | | |
|---|---|---|---|---|---|
| | AIC | BIC | Log Likelihood | Deviance | |
| | 13033.4 | 13070.7 | -6511.7 | 13023.4 | |
| Scaled Residuals: | | | | | |
| | Min | 1Q | Median | 3Q | Max |
| | -4.573 | 0.295 | 0.469 | 0.560 | 0.820 |
| Random Effects: | | | | | |
| | Groups | Name | Variance | Std. Dev | |
| | ID | (Intercept) | 3.670 | 1.916 | |
| | | target_contrast | 7.781 | 2.789 | |
| Number of observations: 12920, groups: ID, 29 | | | | | |
| Fixed Effects | | | | | |
| | | Estimate | Std. Error | z value | |
| | (Intercept) | -0.704 | 0.414 | -1.701 | |
| | target_contrast | 2.731 | 0.585 | 4.671 | |

**Testing for an Effect of S1 Target Contrast (i.e., Discriminability) on Confidence Ratings**

For analyses of confidence ratings, we excluded trials in which the S2 target appeared, as the S2 target biased confidence ratings toward the extreme ends of the scale. We fit linear mixed effects regression models (Gaussian family). We fit separate models for trials with correct responses and for trials with errors.

We compared the following models including effects of S1 target contrast:

```
confidence_normalised ~ target_contrast + (1 | ID)
```

With comparison models that did not include a fixed effect of S1 target contrast:

```
confidence_normalised ~ (1 | ID)
```

Coefficients for the linear mixed effects model (Gaussian family) fit by maximum likelihood to trials with correct responses

| | | | | | |
|---|---|---|---|---|---|
| | AIC | BIC | Log Likelihood | Deviance | |
| | -683.6 | -657.4 | 345.8 | -691.6 | |
| Scaled Residuals: | | | | | |
| | Min | 1Q | Median | 3Q | Max |
| | -4.557 | -0.258 | 0.293 | 0.578 | 1.757 |
| Random Effects: | | | | | |
| | Groups | Name | Variance | Std. Dev | |
| | ID | (Intercept) | 0.010 | 0.010 | |
| | Residual | | 0.050 | 0.223 | |
| Number of observations: 5093, groups: ID, 29 | | | | | |
| Fixed Effects | | | | | |
| | | Estimate | Std. Error | t value | |
| | (Intercept) | 0.383 | 0.031 | 12.20 | |
| | target_contrast | 0.552 | 0.031 | 17.76 | |

Coefficients for the linear mixed effects model (Gaussian family) fit by maximum likelihood to trials with errors

| | | | | | |
|---|---|---|---|---|---|
| | AIC | BIC | Log Likelihood | Deviance | |
| | 834.5 | 855.4 | -413.3 | 826.5 | |
| Scaled Residuals: | | | | | |
| | Min | 1Q | Median | 3Q | Max |
| | -2.284 | -0.707 | 0.089 | 0.804 | 2.617 |
| Random Effects: | | | | | |
| | Groups | Name | Variance | Std. Dev | |
| | ID | (Intercept) | 0.025 | 0.159 | |
| | Residual | | 0.102 | 0.319 | |
| Number of observations: 1366, groups: ID, 29 | | | | | |
| Fixed Effects | | | | | |
| | | Estimate | Std. Error | t value | |
| | (Intercept) | 0.944 | 0.072 | 13.06 | |
| | target_contrast | -0.563 | 0.082 | -6.81 | |

**Supplementary Table S1.** *Summary statistics for numbers of epochs with correct responses retained for analyses per participant, split by condition.*

| Comparison | Condition | Mean Epochs | Median Epochs | SD Epochs | Min Epochs | Max Epochs |
|---|---|---|---|---|---|---|
| Staircase Number | Staircase 1 | 94 | 97 | 11 | 60 | 111 |
| | Staircase 2 | 111 | 114 | 13 | 82 | 129 |
| | Staircase 3 | 115 | 115 | 17 | 84 | 146 |
| By S2 Appears / Absent and Staircase Number | S2 Appears Staircase 1 | 47 | 46 | 6 | 29 | 55 |
| | S2 Appears Staircase 2 | 55 | 56 | 8 | 35 | 69 |
| | S2 Appears Staircase 3 | 58 | 60 | 9 | 41 | 75 |
| | S2 Absent Staircase 1 | 48 | 48 | 7 | 31 | 61 |
| | S2 Absent Staircase 2 | 56 | 56 | 7 | 37 | 65 |
| | S2 Absent Staircase 3 | 57 | 56 | 9 | 40 | 71 |

**Supplementary Table S2.** *Summary statistics for numbers of epochs with erroneous responses retained for analyses per participant, split by condition.*

| Comparison | Condition | Mean Epochs | Median Epochs | SD Epochs | Min Epochs | Max Epochs |
|---|---|---|---|---|---|---|
| Staircase Number | Staircase 1 | 36 | 36 | 11 | 12 | 57 |
| | Staircase 2 | 26 | 21 | 11 | 13 | 53 |
| | Staircase 3 | 24 | 23 | 12 | 6 | 47 |
| By S2 Appears / Absent and Staircase Number | S2 Appears Staircase 1 | 19 | 18 | 6 | 8 | 30 |
| | S2 Appears Staircase 2 | 12 | 10 | 6 | 5 | 30 |
| | S2 Appears Staircase 3 | 12 | 11 | 6 | 1 | 22 |
| | S2 Absent Staircase 1 | 18 | 17 | 6 | 4 | 29 |
| | S2 Absent Staircase 2 | 13 | 12 | 6 | 6 | 27 |
| | S2 Absent Staircase 3 | 12 | 12 | 7 | 3 | 26 |

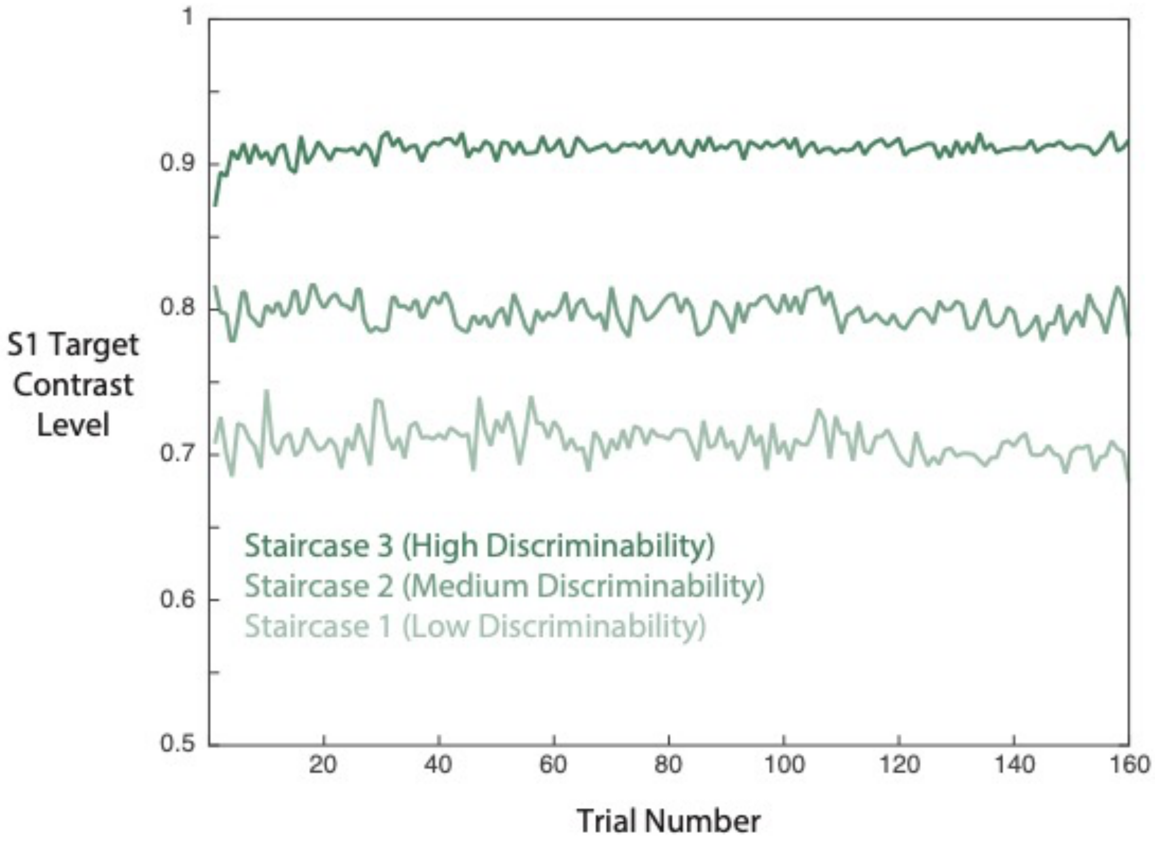


**Supplementary Figure S1.** Group mean S1 target contrast levels by trial for each of the three staircase conditions. Each staircase was used in 160 trials per participant. Trials are ordered from the earliest to the latest within the experiment.

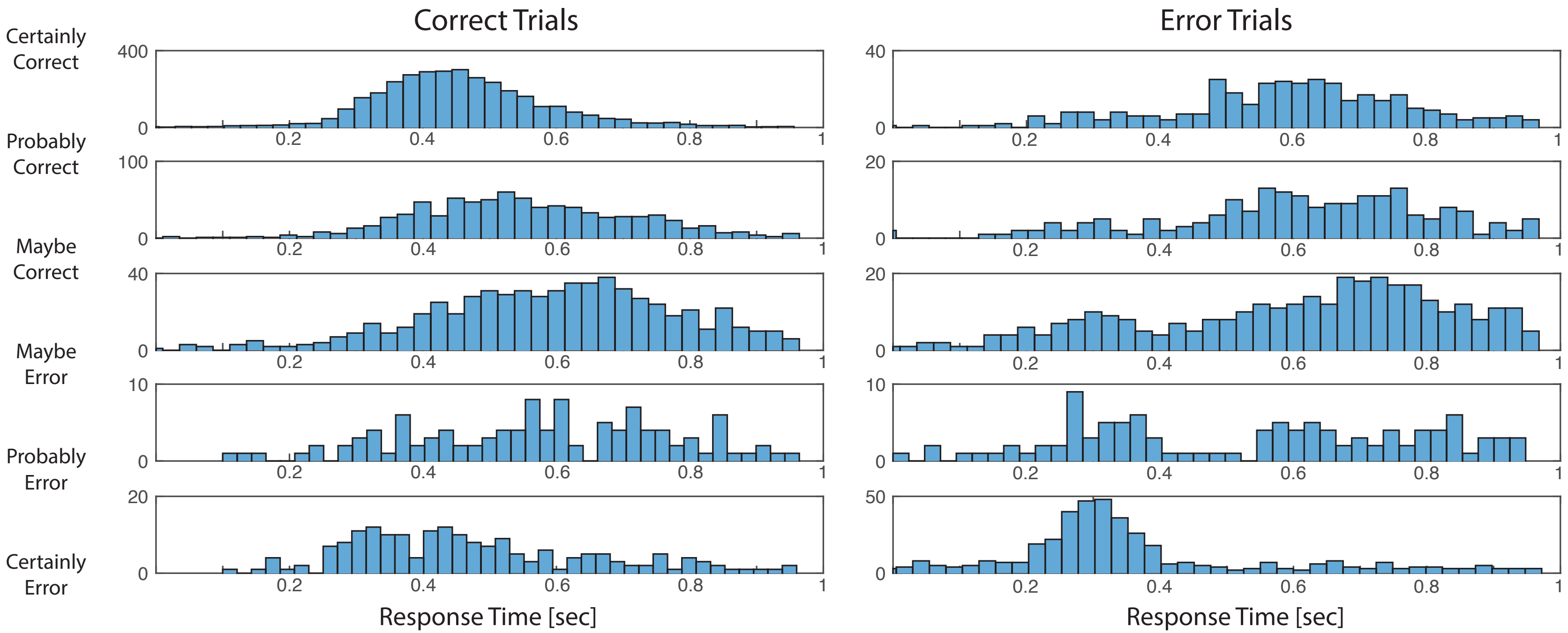


**Supplementary Figure S2**. Histograms of response times (pooled across participants) for different confidence rating bands, for trials with correct responses (left) and errors (right). The confidence indicator labels were used to derive boundaries of each confidence rating band. For example, the top plots show the RT distributions for confidence ratings between 'probably correct' and 'certainly correct'.

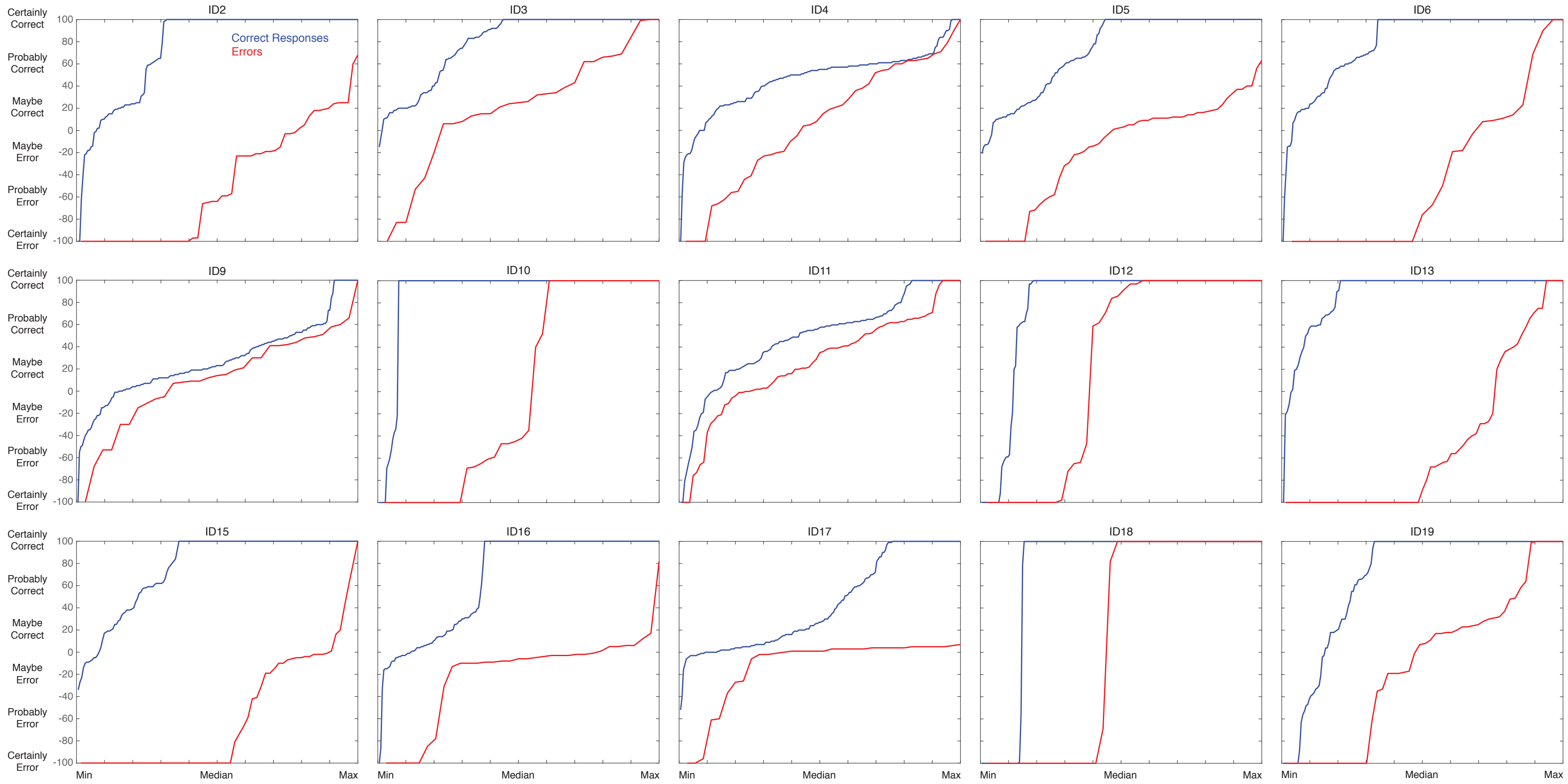


**Supplementary Figure S3**. Confidence rating distributions (ordered from lowest to highest) for each participant. Confidence ratings are plotted separately for trials with correct responses (blue) and errors (red). Data from trials where the S2 target appeared are not included here because the appearance of the easily-discriminable S2 target shifted confidence ratings to the ends of the rating scale. This figure continues on the next page.

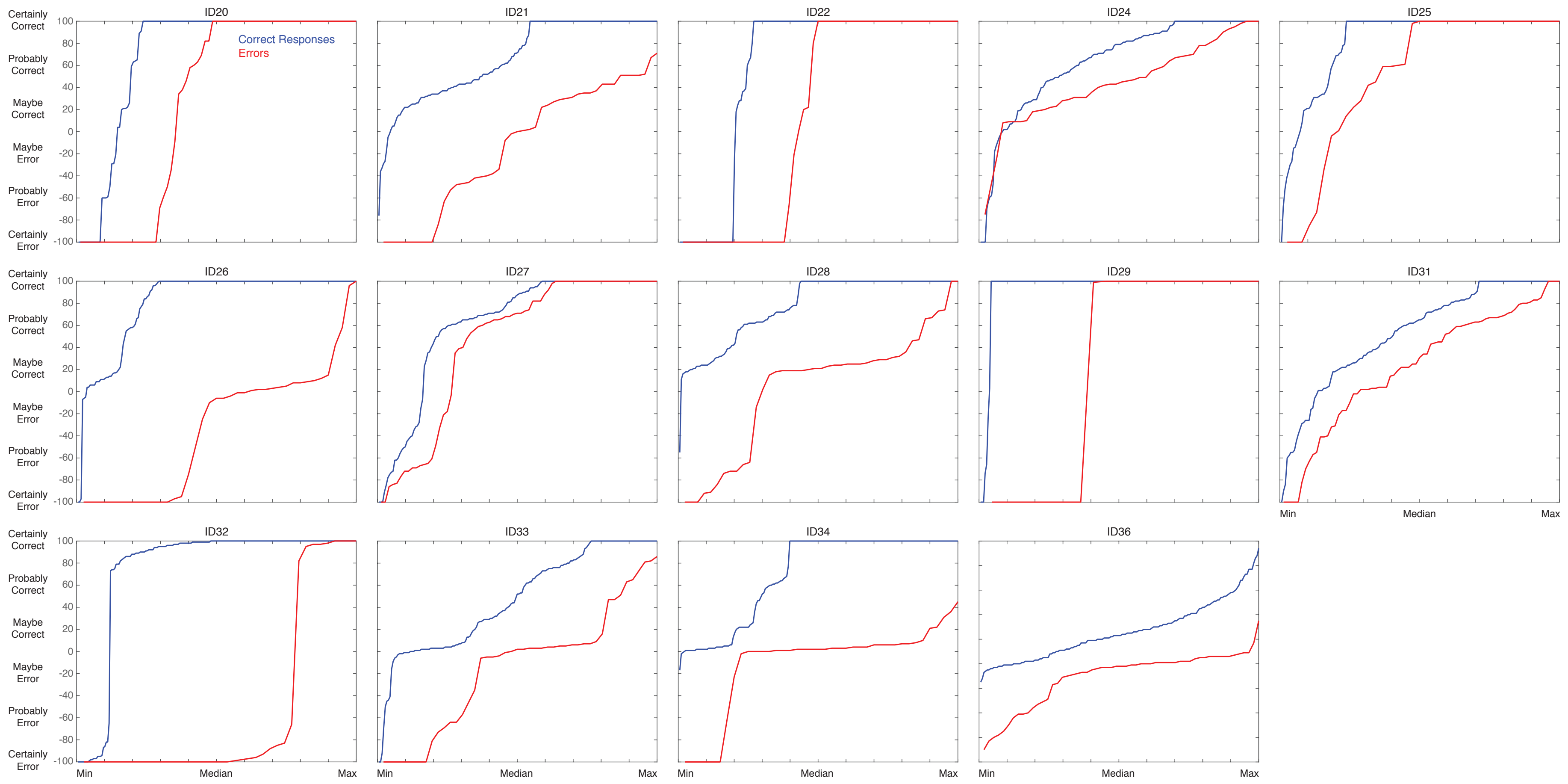


**Supplementary Figure S3 (Continued).**

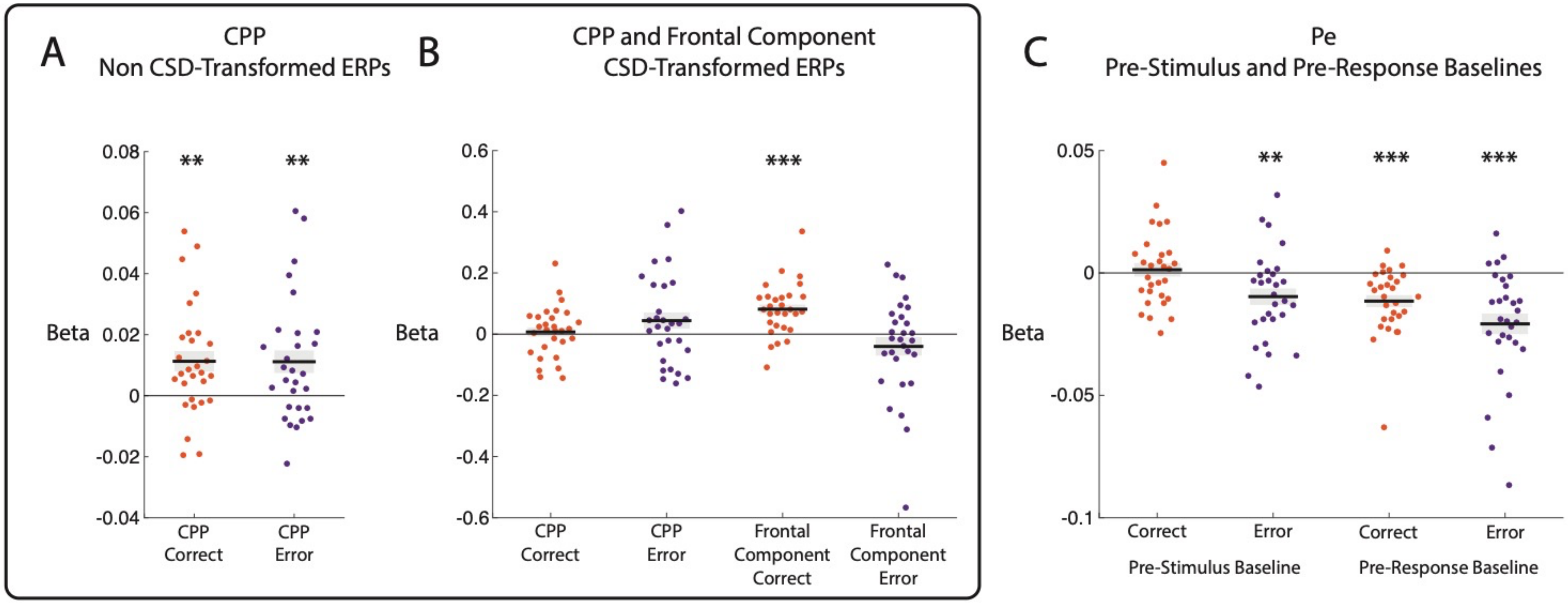


**Supplementary Figure S4.** Results of regression analyses predicting CPP, Frontal Component and Pe mean amplitudes based on confidence ratings in each trial. Beta values correspond to the slopes of linear regression models fit to data for each participant separately. Values above zero signify positive-going associations between ERP component amplitudes and confidence ratings. Values below zero denote negative-going associations. A) Beta values for regressions predicting CPP amplitudes using conventional (i.e., non CSD-transformed) ERPs. B) Beta values for regressions predicting CPP and Frontal Component amplitudes using CSD-transformed ERPs. C) Beta values for regressions predicting Pe amplitudes, for analyses of ERPs baseline-corrected using pre-stimulus and pre-response baselines. Data are shown for correct and error trials separately. Thick black lines represent group mean beta values. Shaded regions denote standard errors. Asterisks denote statistically significant differences from zero at the group level (** denotes $p < .01$ and *** denotes $p < .001$).

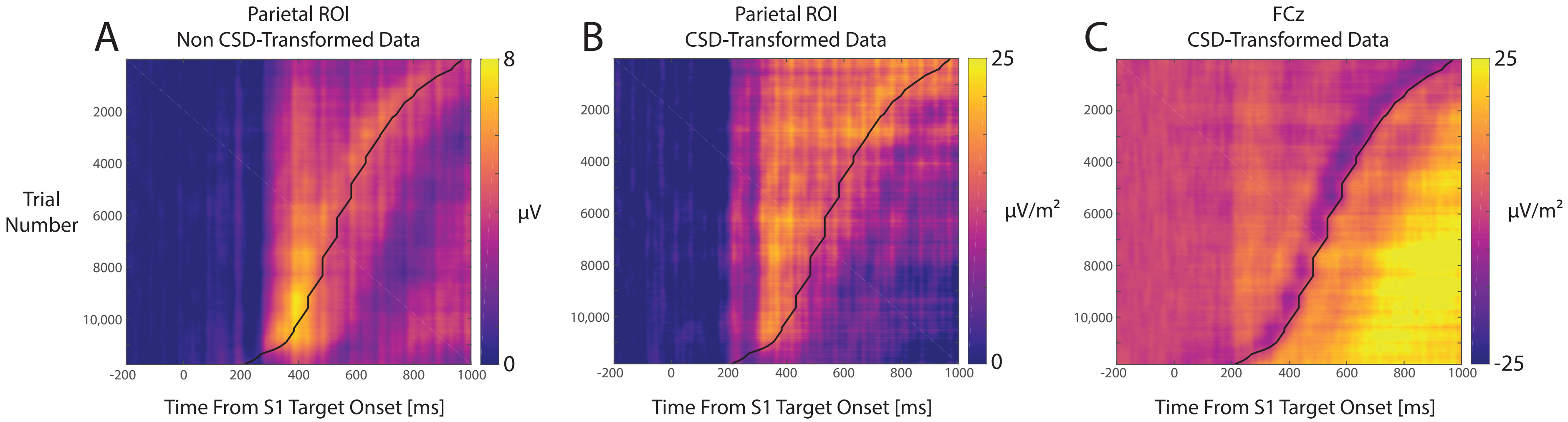


**Supplementary Figure S5**. Heat maps of single trial ERP amplitudes (locked to S1 target onset) pooled across participants and sorted by RT. A.) ERPs at parietal electrodes Pz, P1, P2, CPz, and POz using non CSD-transformed data. B.) ERPs at parietal electrodes Pz and CPz using CSD-transformed data. C.) ERPs at frontal electrode FCz using CSD-transformed data. The curved black lines depict RTs, adjusted to the timing of the following target contrast refresh (to be consistent with how the response-locked ERPs were generated in this study). These plots show clear evidence for response-locked positive-going components (i.e., the CPP) at parietal electrodes, and a response-locked ERP component at FCz. There is also evidence of a stimulus-locked positive-going component at parietal electrodes around 400 ms from S1 target onset. Trials with both correct responses and errors, and trials where the S2 stimulus did and did not appear are included in these plots. ERP amplitudes and RTs were smoothed across trials using a boxcar function spanning 800 trials.

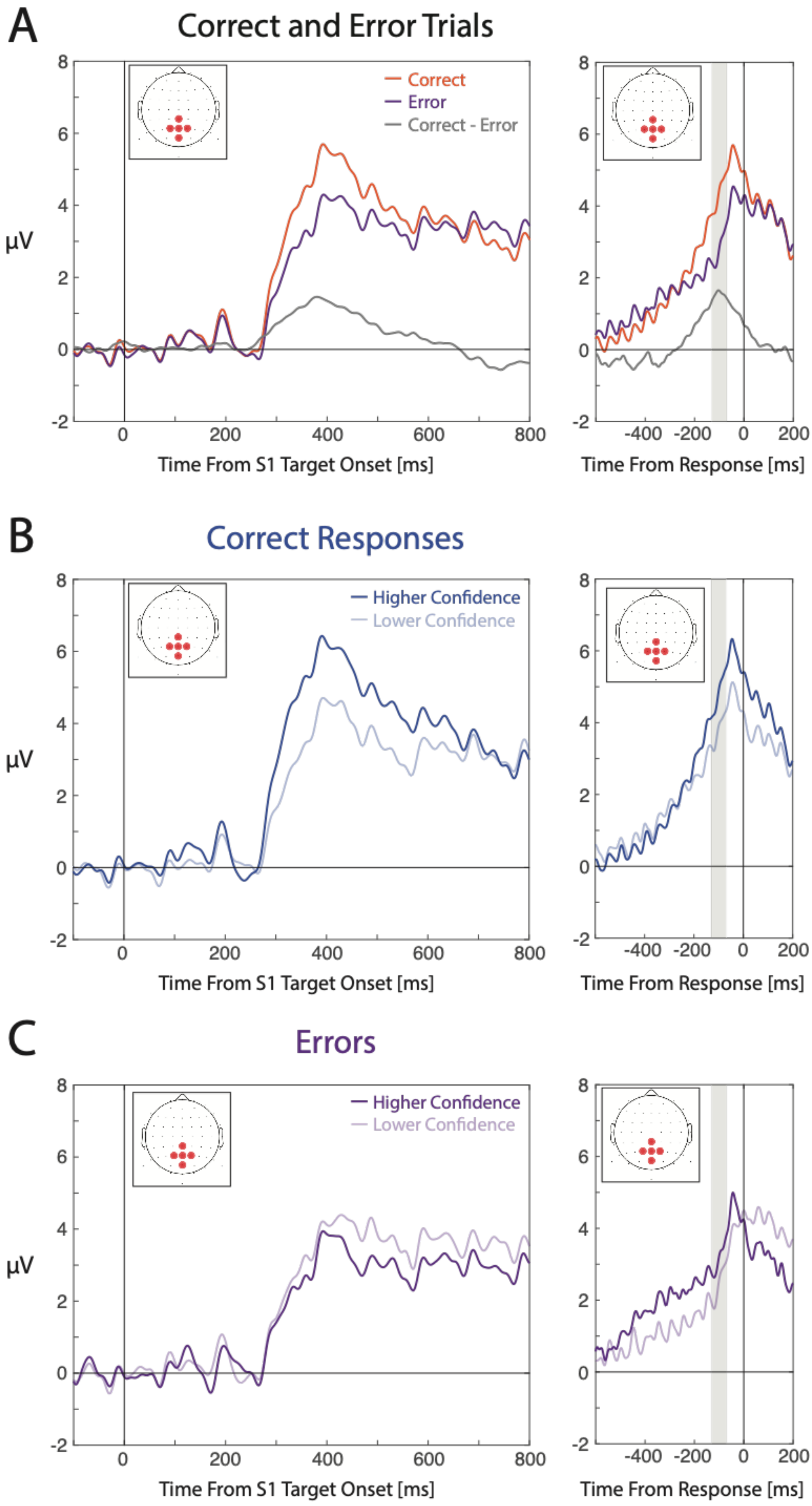
A
Correct and Error Trials
Correct
Error
Correct - Error
μV
Time From S1 Target Onset [ms]
Time From Response [ms]
B
Correct Responses
Higher Confidence
Lower Confidence
μV
Time From S1 Target Onset [ms]
Time From Response [ms]
C
Errors
Higher Confidence
Lower Confidence
μV
Time From S1 Target Onset [ms]
Time From Response [ms]

**Supplementary Figure S6**. ERPs time-locked to S1 target onset (left panels) and the S1 response (right panels) at channels Pz, P1, P2, CPz, and POz. A) ERPs for correct and error responses. Grey shaded regions denote the mean amplitude measurement time windows for the CPP. B) ERPs for higher/lower confidence ratings in trials with correct responses. C) ERPs for higher/lower confidence ratings in trials with errors.

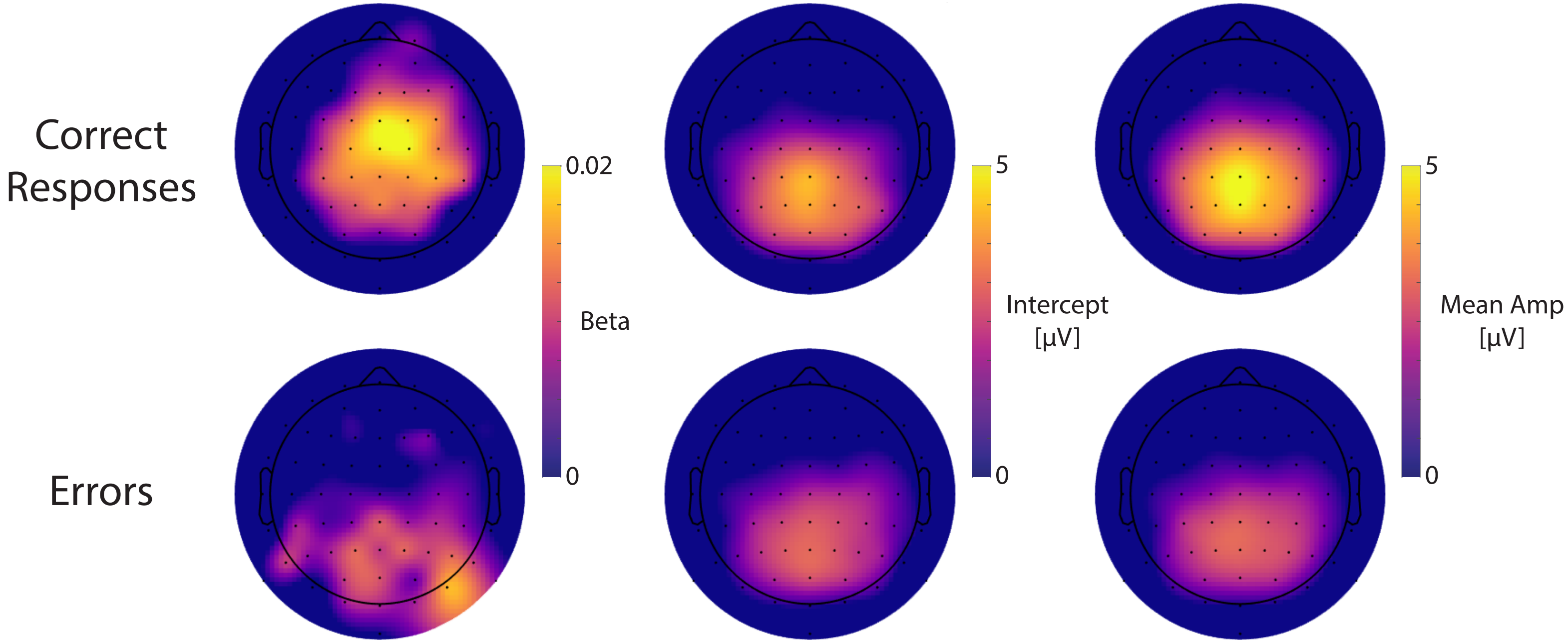


**Supplementary Figure S7**. Scalp maps of group-averaged beta values and intercepts from analyses using confidence ratings to predict ERP amplitudes during the CPP pre-response time window. Separate scalp maps are shown for trials with correct responses (top row) and errors (bottom row). Mean amplitudes during the CPP pre-response time window are shown in the rightmost scalp maps. For correct responses, there is a clear dissociation between the topography of beta values (indicating positive-going associations between confidence ratings and ERP amplitudes) and the topographies of intercepts and mean amplitudes (depicting the typical topography of the CPP in non CSD-transformed data).

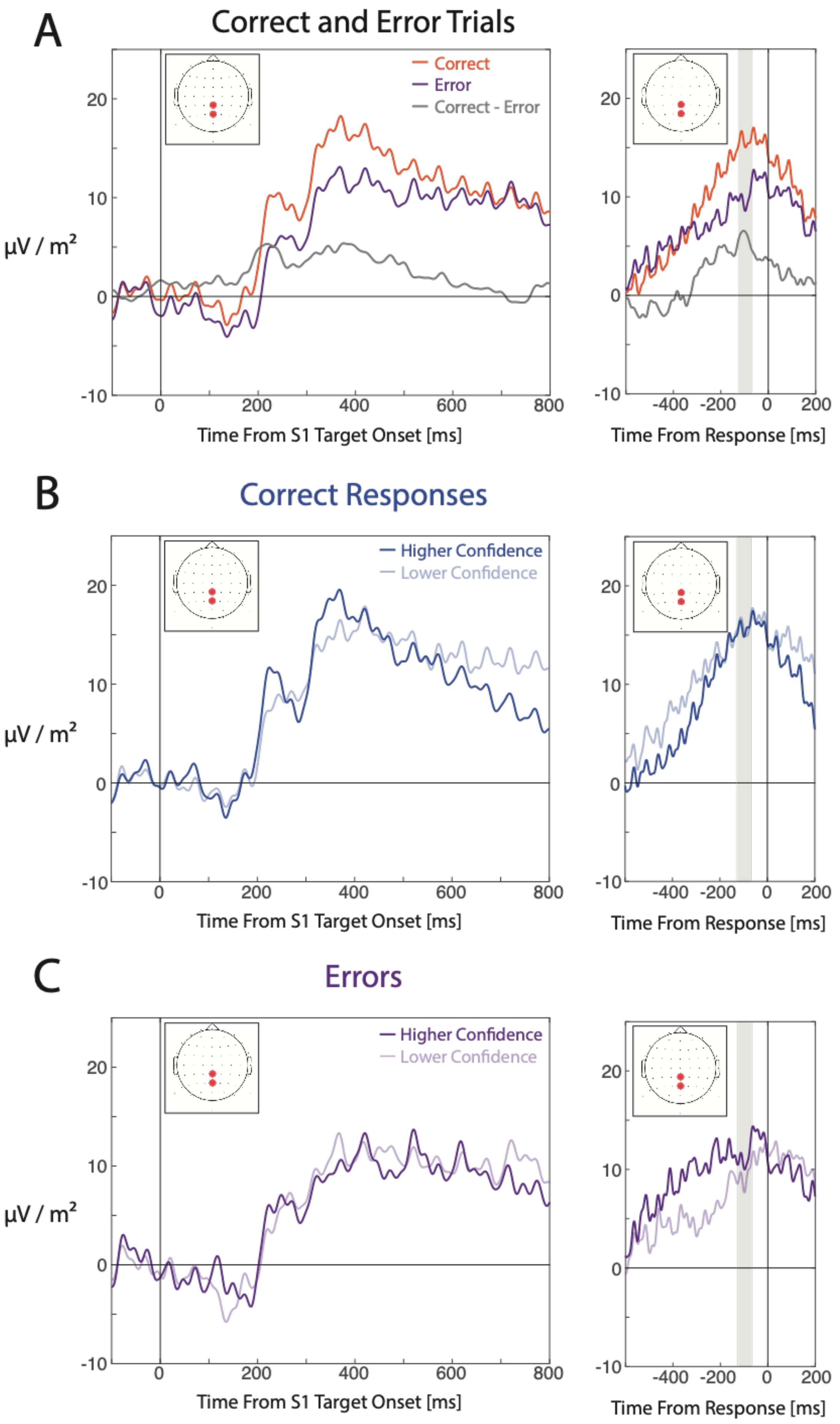
A
Correct and Error Trials
Correct
Error
Correct - Error
µV / m²
20
10
0
-10
0
200
400
600
800
Time From S1 Target Onset [ms]
-400
-200
0
200
Time From Response [ms]
B
Correct Responses
Higher Confidence
Lower Confidence
µV / m²
Time From S1 Target Onset [ms]
Time From Response [ms]
C
Errors
Higher Confidence
Lower Confidence
µV / m²
Time From S1 Target Onset [ms]
Time From Response [ms]

**Supplementary Figure S8**. CSD-ERPs time-locked to S1 target onset (left panels) and the response to the S1 target (right panels) at parietal channels Pz and CPz. A) ERPs for correct and error responses. Grey shaded regions denote the mean amplitude measurement time windows for the CPP. B) ERPs for higher/lower confidence ratings in trials with correct responses. C) ERPs for higher/lower confidence ratings in trials with errors.

### Analyses of ERP Component Amplitudes Using Restricted Confidence Ranges

In addition to the analyses described in the paper, we also used regression models to predict CPP, Frontal Component and Pe amplitudes using restricted ranges of confidence ratings. These included analyses focused on participants' certainty that they had made a correct response, including trials with confidence ratings ranging from "unsure" (0) to "certainly correct" (100). We also ran analyses focused on participants' certainty that they had committed an error, including trials with confidence ratings ranging from "certainly wrong" (-100) to "unsure" (0). The analyses described here included both trials with correct responses and errors. Regression-based analyses were conducted in the same way as described in the paper. Please note that, for the analyses described below, positive-going associations denote instances where more positive-going ERP amplitudes occurred with higher confidence ratings. This means that, for analyses examining certainty in having committed an error, a positive-going association means that more positive-going amplitudes were observed for confidence ratings closer to "unsure" as opposed to "certainly wrong".

#### CPP Component

##### *Non CSD-Transformed ERPs*

CPP pre-response amplitudes were positively associated with confidence across the range of "unsure" to "certainly correct" ($t(28) = 5.08$, $p < .001$, $BF_{10} > 1{,}000$) but not across the range of "certainly wrong" to "unsure" ($t(28) = 0.84$, $p = .406$, $BF_{10} = 0.27$).

##### *CSD-Transformed ERPs*

For CSD-transformed data, CPP pre-response amplitudes were not associated with confidence across the range of "unsure" to "certainly correct" ($t(28) = 1.12$, $p = .273$, $BF_{10} = 0.35$) or across the range of "certainly wrong" to "unsure" ($t(28) = -0.74$, $p = .466$, $BF_{10} = 0.25$).

#### Frontal Component

Frontal Component amplitudes were positively associated with confidence across the range of “unsure” to “certainly correct” ($t(28) = 3.68$, $p < .001$, $BF_{10} = 34.18$) and also negatively associated with confidence across the range of “certainly wrong” to “unsure” ($t(28) = -2.13$, $p = .042$, $BF_{10} = 1.38$). In the latter analysis, the Bayes factor did not indicate strong support for the alternative hypothesis.

### Pe Component

#### *Using Pre-Stimulus Baselines*

When using a pre-stimulus baseline, Pe amplitudes were not associated with confidence across the range of “unsure” to “certainly correct” ($t(28) = 0.06$, $p = .950$, $BF_{10} = 0.20$). Pe amplitudes were negatively associated with confidence across the range of “certainly wrong” to “unsure” ($t(28) = -4.57$, $p < .001$, $BF_{10} > 280$).

#### *Using Pre-Response Baselines*

When using a pre-response baseline, Pe amplitudes were negatively associated with confidence across the range of “unsure” to “certainly correct” ($t(28) = -3.31$, $p = .003$, $BF_{10} = 14.55$) and also across the range of “certainly wrong” to “unsure” ($t(28) = -4.38$, $p < .001$, $BF_{10} > 180$).

### Supplementary Material References

Bates, D., Maechler, M., Bolker, B., & Walker, S. (2015). Fitting linear mixed-effects models using lme4. *Journal of Statistical Software, 67*(1), 1-48.
